# Explosive cytotoxicity of ‘ruptoblasts’ bridges hormonal surveillance and immune defense

**DOI:** 10.1101/2025.03.28.645876

**Authors:** Chew Chai, Eliya Sultan, Souradeep R. Sarkar, Lihan Zhong, Dania Nanes Sarfati, Orly Gershoni-Yahalom, Christine Jacobs-Wagner, Hawa Racine Thiam, Benyamin Rosental, Bo Wang

## Abstract

Current understanding of cytotoxic immunity is shaped by hematopoietic-derived cells – T cells, natural killer cells, and neutrophils. Here, we identify ‘ruptoblasts’, a previously unknown cytotoxic glandular cell type in regenerative planarian flatworms. Ruptoblasts undergo an explosive cell death, ‘ruptosis’, triggered by activin, a multifunctional hormone that also acts as an inflammatory cytokine. Excessive activin – induced through protein injection, genetic chimerism, or bacterial infection – initiates ruptosis, discharging potent diffusible cytotoxic agents capable of eliminating any nearby cells, bacteria, and even mammalian cells within minutes. Ruptoblast ablation suppresses inflammation but compromises bacterial clearance, highlighting their broad-spectrum immune functions. Mechanistically distinct from known cytotoxic mechanisms, the explosive nature of ruptosis relies on intracellular calcium and dynamic cytoskeletal reorganization. Ruptoblast-like cells appear conserved in diverse basal bilaterians, implying an ancient evolutionary origin. These findings unveil a widespread strategy coupling hormonal regulation with immune defense and expand the landscape of evolutionary immune innovations.

## Introduction

Cytotoxicity is a core immune function essential for eliminating pathogens, aberrant cells, and malignant clones. In animals, major cytotoxic cells include cytotoxic T cells and natural killer cells, which kill via direct cell-to-cell contact, and neutrophils, which deploy diffusible agents such as reactive oxygen species, proteases, and extracellular DNA traps^1–4^. Despite differences in their killing mechanisms, all known cytotoxic immune cell types share a common evolutionary origin from the hematopoietic lineage^5–9^, raising the question of whether cytotoxicity is inherently restricted to this lineage and whether other cell types can independently acquire cytotoxic function through distinct mechanisms. Exploring alternative origins of cytotoxicity could reveal fundamental principles underlying immune evolution and offer new strategies for therapeutic intervention.

As a potential context for discovering novel cytotoxic cells, we focused on a function specific to adaptive immunity. In mammals, cytotoxic T cells can recognize autoantigens produced during hormone secretion, such as pre-proinsulin (precursor of insulin) and thyroglobulin (precursor of thyroid hormones), and selectively eliminate hypersecreting cells (**Figure 1A**)^10^. This function is particularly crucial in proliferative endocrine tissues, such as the pancreas, thyroid, and adrenal gland, where continuous cell divisions increase the risk of mutations that can lead to hypersecreting clones^11^. Dysregulation of this immune surveillance mechanism can lead to autoimmune disorders such as type I diabetes and Hashimoto’s thyroiditis^12, 13^. We reasoned that, if basal animals without adaptive immunity possess a similar function, it should be mediated through an entirely novel cytotoxic cell lineage.

**Figure 1.**
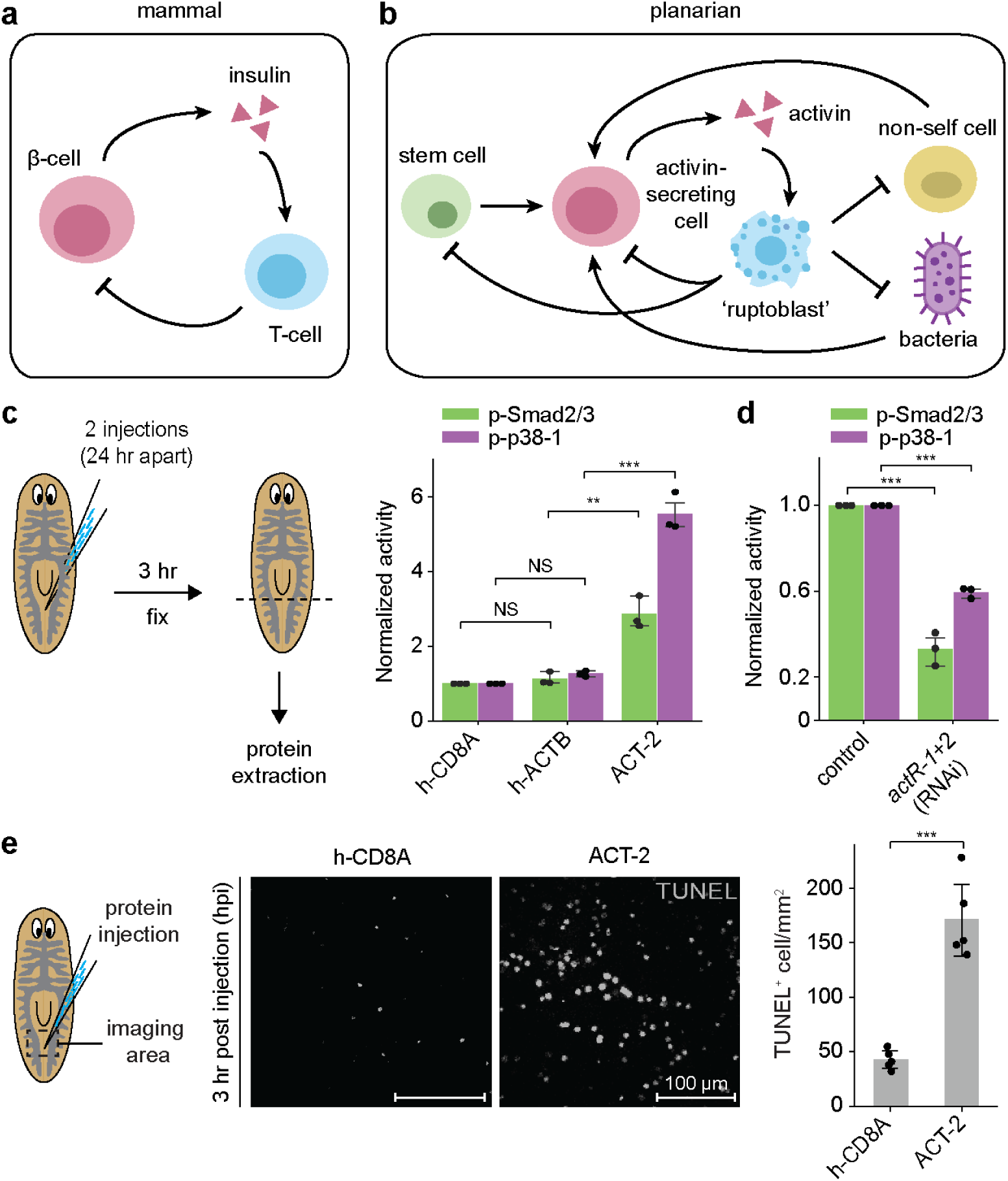
ACT-2 induces inflammatory responses in planarians. **(A)** Schematics showing T cell-mediated elimination of hormone-hypersecreting cells in mammals, using insulin-producing *β*-cells as an example. **(B)** Proposed activin surveillance mechanism in planarians in which overactivation of activin signaling causes ‘ruptoblasts’ to eliminate any nearby cells including stem cells, activin-secreting cells, non-self cells and microbes. **(C)** Quantification of p-Smad2/3 (green) and p-p38-1 (magenta) levels using Western blotting in control animals (injected with h-CD8A, h-ACTB) and animals injected with ACT-2 into the parenchymal tissue between posterior gut branches. Tail tissues (posterior to the dashed line) were collected for protein extraction. **(D)** ACT-2 injection-induced increase of p-Smad2/3 and p-p38-1 is reduced by RNAi-mediated knockdown of *actR1* and *actR2*, which are presumptive receptors for ACT-2 (Figure S2E). **(E)** (Left) Representative image of TUNEL staining at the injection site showing cell death at 3 hours post injection (hpi) in animals injected with h-CD8A and ACT-2 proteins, respectively. Dashed boxes: imaging area. (Right) Number denistiy of TUNEL^+^ cells near the injection site. Data points in (C-D) represent biological replicates each containing tail fragments from five animals, and in (E) individual animals pooled from two independent experiments. Statistical significance is determined by a two-sided t-test, with error bars denoting standard deviation (SD). **p < 0.01, ***p <0.001, NS, not significant.

To explore this possibility, we studied the highly regenerative planarian flatworm, *Schmidtea mediterranea*, as its tissues contain abundant proliferative somatic stem cells known as neoblasts^14–16^. These neoblasts divide every couple of days to maintain tissue turnover and support regeneration, thus posing similar risks of somatic mutations and hormonal imbalances as seen in mammalian proliferative organs^17^. However, unlike mammalian T cells, which predominantly target differentiated hormone-secreting cell types (such as β-cells^10, 18^) that produce autoantigens, any immune surveillance mechanism in planarians must also eliminate mutated neoblasts, because otherwise they could replenish aberrant hormone-secreting cells, perpetuating the imbalance.

In planarians, regeneration, reproduction, and tissue homeostasis are all regulated by activin, a hormone whose level must be finely balanced and stably maintained: excessive activin can impair regeneration^19–21^, whereas insufficient levels hinder both asexual^22^ and sexual reproduction (**Figure S1**). Here, we discovered that activin also functions as a potent inflammatory cytokine in response to genetic chimerism and bacterial pathogens, directly linking hormonal regulation with immune responses. Elevated activin levels trigger a previously undescribed immune cell type, ‘ruptoblasts’, to undergo ‘ruptosis’, an explosive cytotoxic event that eradicates nearby cells. This broad-acting cytotoxicity can simultaneously eliminate activin-secreting cells, neoblasts, bacterial pathogens in planarians (**Figure 1B**), and notably, also kill cells from other species (e.g., mammalian cells). In contrast to conventional immune recognition mechanisms that rely on pathogen- or damage-associated cues, ruptoblasts are activated by localized activin signaling, which confers both activation and spatial specificity.

Our analyses also suggest that ruptoblast-like cells, which lack canonical immune markers, belong to a glandular/secretory cell type family^23^. These cells appear evolutionarily ancient, conserved from the base of the bilaterian tree, yet curiously absent from common model organisms. Although ruptosis involves the release of granular contents in a manner reminiscent of neutrophil and mast cell degranulation^3, 4^, its killing mechanism is distinct from any previously described cytotoxic or cell death processes (**Table S1**). Our findings thus uncover a previously unrecognized immune cell type with broad-spectrum activity, highlighting evolutionary convergence in immune strategies through entirely different cellular origins and molecular mechanisms.

## Results

### Excessive activin induces inflammatory responses in planarians

Activin is a pleiotropic hormone that regulates diverse physiological processes in planarians. To test whether excess activin is sufficient to trigger inflammation, we generated recombinant proteins for ACT-2, one of the two planarian activin homologs^19^. We injected planarians (CIW4 asexual clonal line) with either ACT-2 or control proteins (human CD8A and Activin-B), and used Western blotting to measure phosphorylated Smad2/3 (p-Smad2/3), the canonical effector of activin signaling. As expected, p-Smad2/3 levels increased only in ACT-2 injected animals (**Figure 1C** and **S2A**), confirming the functionality and specificity of our recombinant proteins.

We then examined whether activin activated the p38 pathway, which was previously shown to drive inflammatory tissue degeneration during bacterial and fungal infection in planarians^25, 26^. Indeed, only in ACT-2 injected planarians did we detect elevated phosphorylation of p38-1 (p-p38-1) (**Figure 1C**, **S2B** and **S2C**). Knockdown of *act-2* or activin receptors, drastically reduced both p-Smad2/3 and p-p38-1 levels during either homeostasis or after ACT-2 injection (**Figure 1D**, **S2D**, **S2E** and **S2F**), further confirming that *p38-1* acts downstream of activin. Phenotypically, ACT-2 induced tissue lesions and excessive cell death at the injection sites, quantified by TUNEL assay, both of which were alleviated by RNA interference (RNAi) against *p38-1* (**Figure 1E** and **S2G**). These data suggest that excess activin triggers a p38-mediated inflammatory response, establishing a direct link between hormonal dysregulation and immune activation.

### Activin functions as an inflammatory cytokine in genetic chimeras

Since ACT-2-induced lesions resolved within 24 hr, presumably due to degradation or clearance of the recombinant proteins^27^, we sought a method to sustain activin overactivation. Drawing inspiration from classic experiments fusing distinct planarian species^28^, we wondered whether this approach could provoke immune rejection analogous to that commonly observed in mammalian organ transplant. We optimized a protocol^29^ to generate genetic chimeras by fusing asexual (A) and sexual (S) *S. mediterranea* along the midline (**Figure 2A**, **S3A** and **S3B**). Despite seemingly complete tissue integration – such as shared anterior and posterior poles and midline, integrated muscle networks, nervous system, gut branches, and a single pharynx (**Figure 2B**, **S3C** and **S3D**) – these genetic chimeras exhibited three key pathological features, indicative of a chronic inflammatory response to incompatible genotypes.

**Figure 2.**
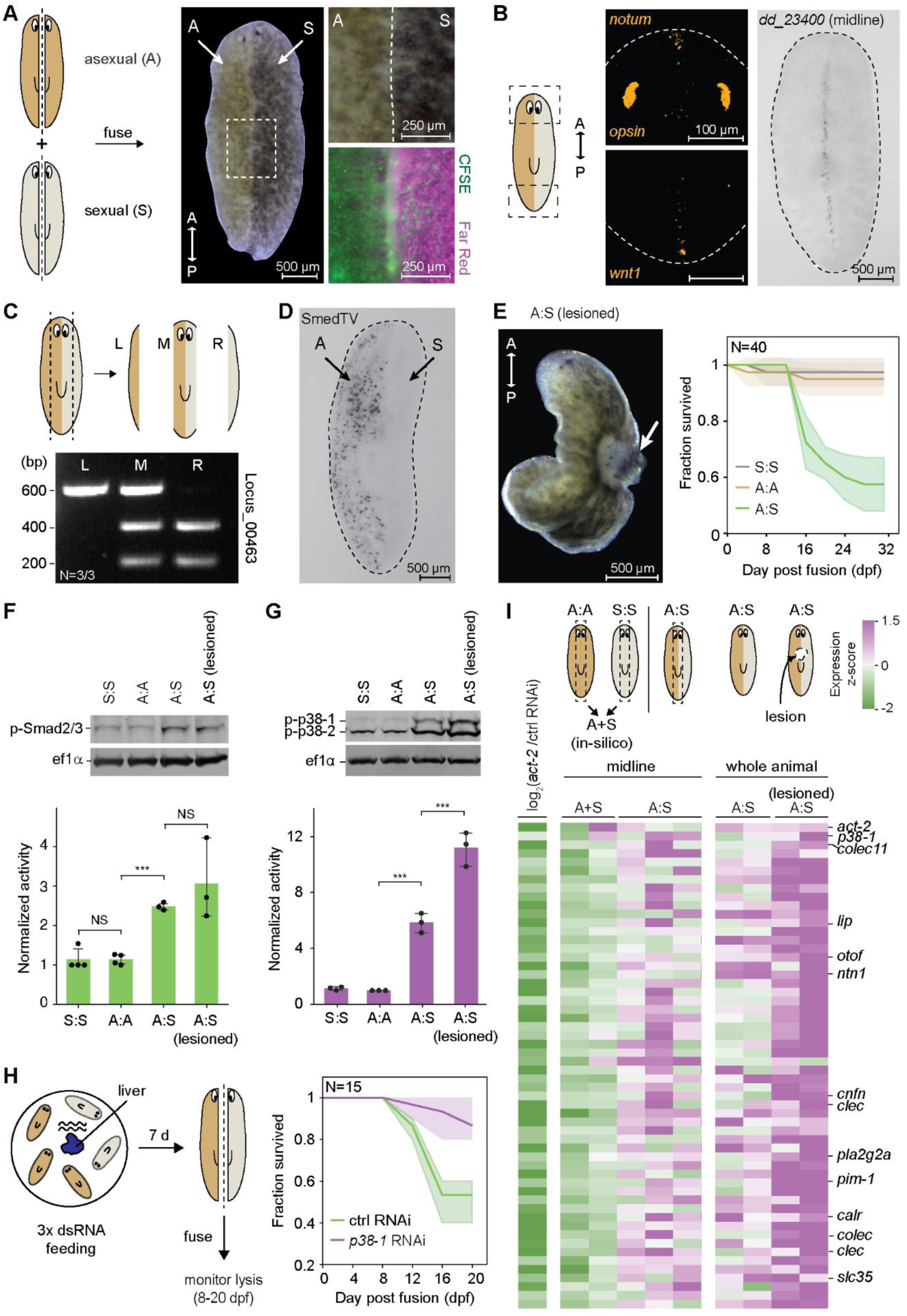
Genetic chimerism induces activin overactivation, resulting in chronic inflammation. **(A)** Planarians of two distinct genotypes, asexual (A) and sexual (S), are cut longitudinally along the midline and fused to form a chimera. Note the different pigmentation of tissues from two genotypes. Right: a magnified view showing the fusion site at 3 days post fusion (dpf) in bright field (top), in fluorescence using tissues pre-stained with CellTrace CFSE and Far Red (bottom). **(B)** (Left) FISH images showing expression of *notum* in the anterior pole and *wnt-1* in the posterior pole of genetic chimeras at 20 dpf. (Right) WISH image showing a single midline marked by *dd_23400* in genetic chimeras at 20 dpf. **(C)** PCR-RFLP analysis of tissues collected from the left (L), middle (M), and right (R) portions of the chimera body. Genotyping is based on locus 00463 cut by ScaI in sexual cells but not in asexual cells. **(D)** WISH image showing the expression of asexual-specific RNA virus SmedTV in genetic chimeras at 20 dpf. Note the clear division between asexual and sexual halves. **(E)** (Left) Brightfield image of a chimera with a lesion at the fusion site. (Right) Survival curves of chimeric and homotypic fusions. **(F-G)** (Top) Western blot showing p-Smad2/3 (F) and p-p38-1(G) in chimeras with lesions at 14– 18 dpf and without lesions at 20 dpf, compared to homotypic fusions. (Bottom) Quantified fold activation normalized to the levels in sexual homotypic fusions. **(H)** (Left) Schematics showing the workflow of RNAi feeding before fusion. (Right) *p38-1* RNAi reduced the fraction of lesioned chimeras. Since *act-2* RNAi induces anteroposterior (AP) axis bifurcation^19^, preventing chimeric fusion. Consequently, this precludes a direct test of whether *act-2* RNAi can mitigate lesion. **(I)** (Left) Heatmap showing expression fold change after *act-2* RNAi (data from Ref. 19). (Middle) Normalized expression comparing midlines of chimeras without lesions to that of numerical average of sexual and asexual homotypic fusions. Genes significantly upregulated (p-value < 0.05, two-sided Welch’s t-test) were shown with *act-2* and *p38-1* added to the top. Genes labeled have immune functions in other organisms. (Right) Comparison of lesioned and non-lesioned chimeras show that the upregulation in chimeras are further amplified during lesion. N in (C) denotes the number of biological replicates that show results consistent with the gel image. SD of survival curves in (E, H) was obtained from three independent experiments with N denoting the total number of animals monitored. Data points in (F, G) represent biological replicates each containing three chimeras. Statistical significance is determined by a two-sided t-test, with error bars denoting SD. ***p <0.001, NS, not significant.

First, the two genotypes remained segregated, as measured by PCR-RFLP genotyping^30^ (**Figure 2C**) and strain-specific RNA virus (SmedTV)^31^ distribution at 20 days post fusion (dpf) (**Figure 2D**). This indicated a rejection between the two genotypes, as neoblasts and/or postmitotic progenitors shouls have significant migratory capacity^32^. Second, unlike homotypic fusions, the chimeras failed to feed, a trait typically associated with sick animals. Finally, ∼40% of chimeras developed lesions at the fusion site by ∼14 dpf, resembling the phenotype induced by ACT-2 protein injections, and then progressed to whole-body disintegration within a day (**Figure 2E**).

To investigate if these lesions stemmed from activin overactivation, we measured p-Smad2/3 (**Figure 2F**) and p-p38-1 (**Figure 2G**) in chimeras and observed markedly increased levels of both. Knocking down *p38-1* prior to fusion prevented these lesions, establishing a causal link between molecular inflammatory responses and the lesion phenotype (**Figure 2H**). We next asked whether inflammation was systemic or confined to the midline, where the two genotypes interacted. RNAseq of both midline tissues and whole chimeras (with and without lesions) revealed that genes downregulated after *act-2* RNAi^19^ were globally upregulated relative to simple averages of homotypic fusions, especially in animals with lesions (**Figure 2I**), suggesting a body-wide inflammatory response. Together, our results demonstrate that genetic chimerism induces chronic inflammation, with activin serving as a inflammatory cytokine.

### Ruptoblasts undergo a unique form of cytotoxic cell death, ruptosis

To identify cells driving the activin-induced inflammatory responses, we dissociated planarians and monitored cellular responses to ACT-2 *ex vivo*. While p-p38-1 level increased within 3 min of ACT-2 treatment in a concentration-dependent manner (**Figure 3A** and **S4A**), we were surprised to observe a subset of cells undergoing explosive lysis within ∼2 min of ACT-2 exposure (**Figure S4B**, **Video S1**). In contrast, exposure to control proteins (h-CD8A, h-ACTB) had minimal effects. Flow cytometry confirmed that activin-induced lysis is dose-dependent (**Figure 3B**), regardless of the planarian genotype (**Figure S4C**). Thus, ACT-2 itself, rather than allogeneic response to non-self, is sufficient to trigger cell lysis.

**Figure 3.**
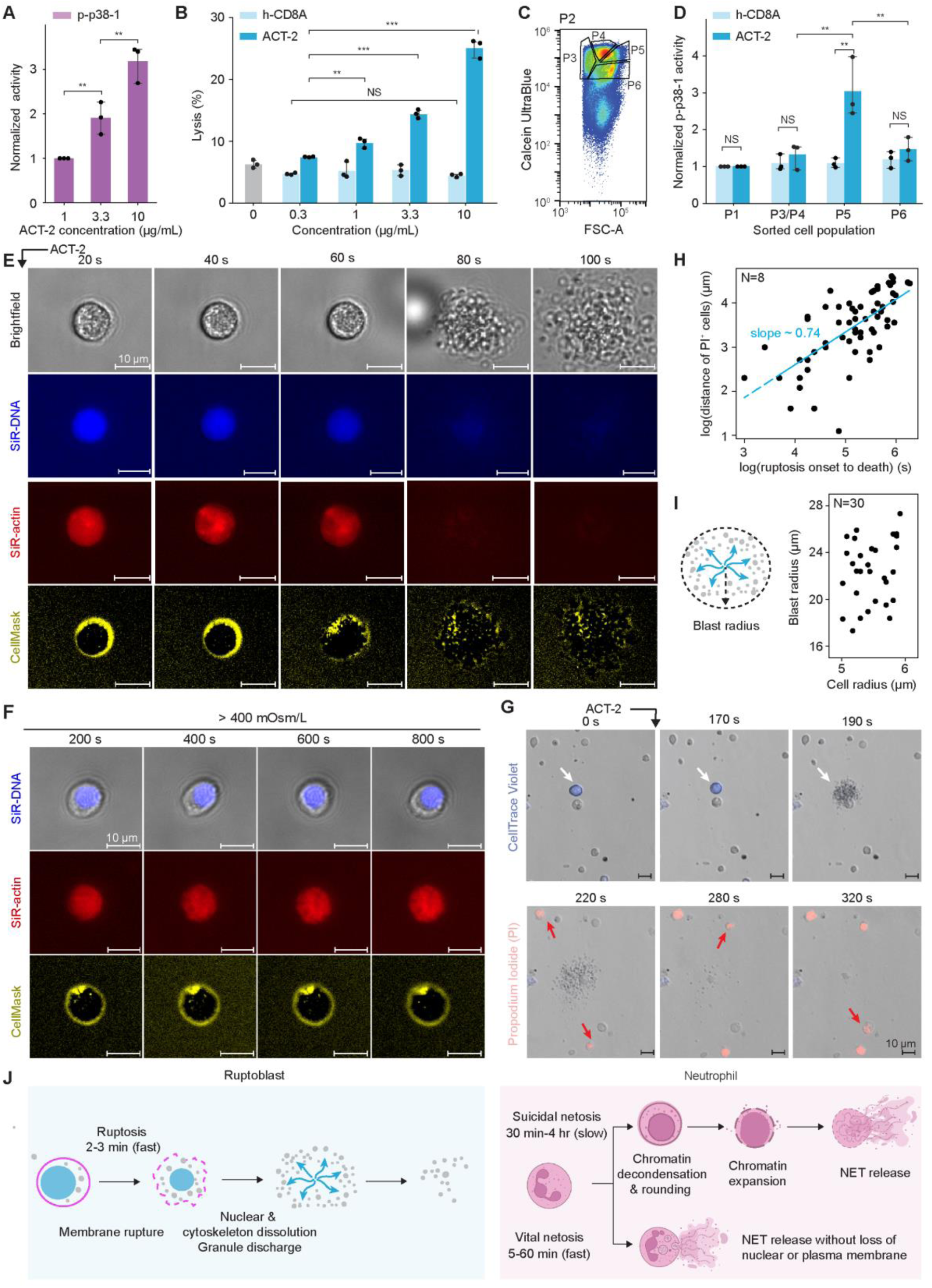
Activin induces ruptosis, an explosive cytotoxic form of cell death. **(A)** Quantification of p-p38-1 in bulk cells following exposure to increasing concentrations of ACT-2. **(A)** Cell death measured via flow cytometry in response to varying concentrations of h-CD8A (control) and ACT-2 proteins, quantified by the percentage of PI^+^ cells. **(C)** Sorting strategy for enriching activin-responding cells based on Calcein Ultrablue stain and forward scattering (FSC), after an initial back scatter (BSC, measuring granularity) gating shown in **Figure S5A**. **(D)** Barplot quantifying p-p38-1 in sorted subpopulations following exposure to 10 μg/mL h-CD8A or ACT-2. P1 is a low back scattering control population (Figure S5A). P3 and P4 are pooled for Western blot to collect sufficient material. **(E)** Snapshots showing key cellular component undergoing complete destruction during ruptosis. Membrane ruptures starting at ∼60 s as nucleus disintegrates and actin disassembles, culminating in the complete breakdown of the entire cell and explosive discharge of granules. **(F)** Snapshots showing ruptoblasts incubated in the calcium/magnesium free (CMF) media with sucrose to elevate external osmolarity. Ruptosis is completely suppressed at 400 mOsm/L, with no nuclear fragmentation (top), cytoskeleton disassembly (middle) and membrane rupture (bottom) or granule discharge, indicating that high external osmotic pressure can prevent the execution of ruptosis. **(G)** Snapshots demonstrating that sorted P5 cells (CellTrace Violet-labeled) can kill nearby unlabeled cells (pooling P1, P3, P4, P6). Cell death is indicated by PI entry (red). Cells were mixed at a 1:4 ratio of P5 to the rest. ACT-2 was added at ∼90 s. White arrow: ruptoblast, red arrows: killed cells. **(H)** Timing of PI uptake of individual cells as a function of distance from the ruptoblast. Line: power law with an exponent of ∼ 0.7, suggesting super-diffusive spread of cytotoxic agents. **(I)** Quantification of maximum dispersal radius of granules released during ruptosis. **(J)** Comparison of ruptosis and NETosis. Ruptosis is a rapid, all-or-none event involving mostly synchronized membrane rupture, actin depolymerization, and DNA fragmentation, culminating in the explosive release of cytotoxic granules. In contrast, some cellular components (DNA for suicidal NETosis and plasma membrane and nucleus for vital NETosis) can persist through NETosis. Schematics for neutrophils are created with BioRender.com. Data points represent biological replicates each consisting of 50,000 cells in (A), and 300,000 cells in (B, D). Statistical significance is determined by a two-sided t-test, with error bars denoting SD from three independent experiments. For (E,F), the corresponding videos are shown in **Video S2**, and for (G), the corresponding video is shown in **Video S3**. **p < 0.01, ***p <0.001, NS, not significant.

We next developed a fluorescence activated cell sorting (FACS) strategy to enrich for these explosive cells. Based on morphology, we gated for high-granularity cells (**Figure S5A**) and subdivided by cell size and a cytoplasmic stain (Calcein Ultrablue) (**Figure 3C**). Testing each subpopulation’s response to ACT-2, we identified a single small (<3% of total cells) population (P5) that activated p38 (**Figure 3D**). A major fraction (∼60-70%) of these cells burst upon activin exposure, releasing numerous granules that dispersed and vanished within 2-5 min (**Figure 3E, S5B**, and **S5C**, **Video S1**). Shortly after plasma membrane (labeled by CellMask) rupture and granule discharge, both the actin cytoskeleton (labeled by SiR-actin) and the nucleus (labeled by SiR-DNA) were completely disintegrated (**Figure 3E**, **Video S2**), indicating rapid cytoskeletal disassembly and DNA cleavage. We refer to this explosive cell death as ‘ruptosis’ and these cells as ‘ruptoblasts’.

The explosive nature of ruptosis indicates an instantaneous and forceful discharge of cellular contents. When we increased extracellular osmolarity by adding sucrose to the media, ruptosis was entirely inhibited at osmolarities above 400 mOsm/L (**Figure 3F**, **S5D**, and **Video S2**), with no evidence of granule discharge, membrane rupture, actin disassembly, or nuclear disruption. This result suggests that ruptosis depends on a net outward force, potentially generated by an imbalance between intracellular and external pressures.

To test whether ruptosis kills surrounding cells, we labeled P5 cells with CellTrace Violet and co-incubated them with unlabeled, P5-depleted cell fractions. Upon inducing ruptosis with ACT-2, all cells near ruptoblasts rapidly took up propidium iodide (PI), indicative of cell death (**Figure 3G, Video S3**). The timing of PI uptake relative to the cell’s distance from the ruptoblast followed a power-law scaling with an exponent of ∼0.7, consistent with a super-diffusive spreading of the cytotoxic agent rather than directed, inertia-driven jet (**Figure 3H**). Additionally, the observed killing radius (∼100 μm) exceeded the maximum dispersal range of granules (∼30 μm) (**Figure 3I**), supporting the conclusion that ruptoblast cytotoxicity is mediated by diffusible agents released upon ruptosis.

The speed and scale of cellular destruction during ruptosis is remarkable compared to other known cell death mechanisms (**Figure 3J**). Suicidal NETosis, the major form of sacrificial cytotoxic cell death, typically unfolds over hours^33, 34^. During NETosis, neutrophil DNA does not degrade but decompacts^33, 35, 36^. While neutrophil/mast cell degranulation and vital NETosis (the release of small extracellular traps without full cell rupture) can occur on a minute timescale as ruptosis^34^, these processes occur without plasma membrane rupture^36^.

### Ruptoblasts are glandular/secretory cells

To determine the molecular identity of ruptoblasts, we performed single-cell RNAseq on sorted P5 cells briefly treated with either control (CD8A) or ACT-2 proteins (**Figure 4A**). Under this condition, we anticipated ruptoblasts to be selectively depleted in the ACT-2-treated sample before widespread killing occurred. Among the 16 transcriptomically distinct populations identified, three cell clusters (11, 13 and 15) were markedly depleted (**Figure 4B** and **S6A**). These three clusters expressed high levels of *actR1* and *p38-1* but not *act-2*, implicating that they only sense and respond to activin without amplifying it via activin secretion (**Figure 4C**). Based on known marker genes^23, 24^ (**Figure 4C**, **S6A** and **S6B**), clusters 11 and 13 were annotated as parenchymal glandular/secretory cells, while cluster 15 were *cathepsin^+^*phagocytic cells^37^.

**Figure 4.**
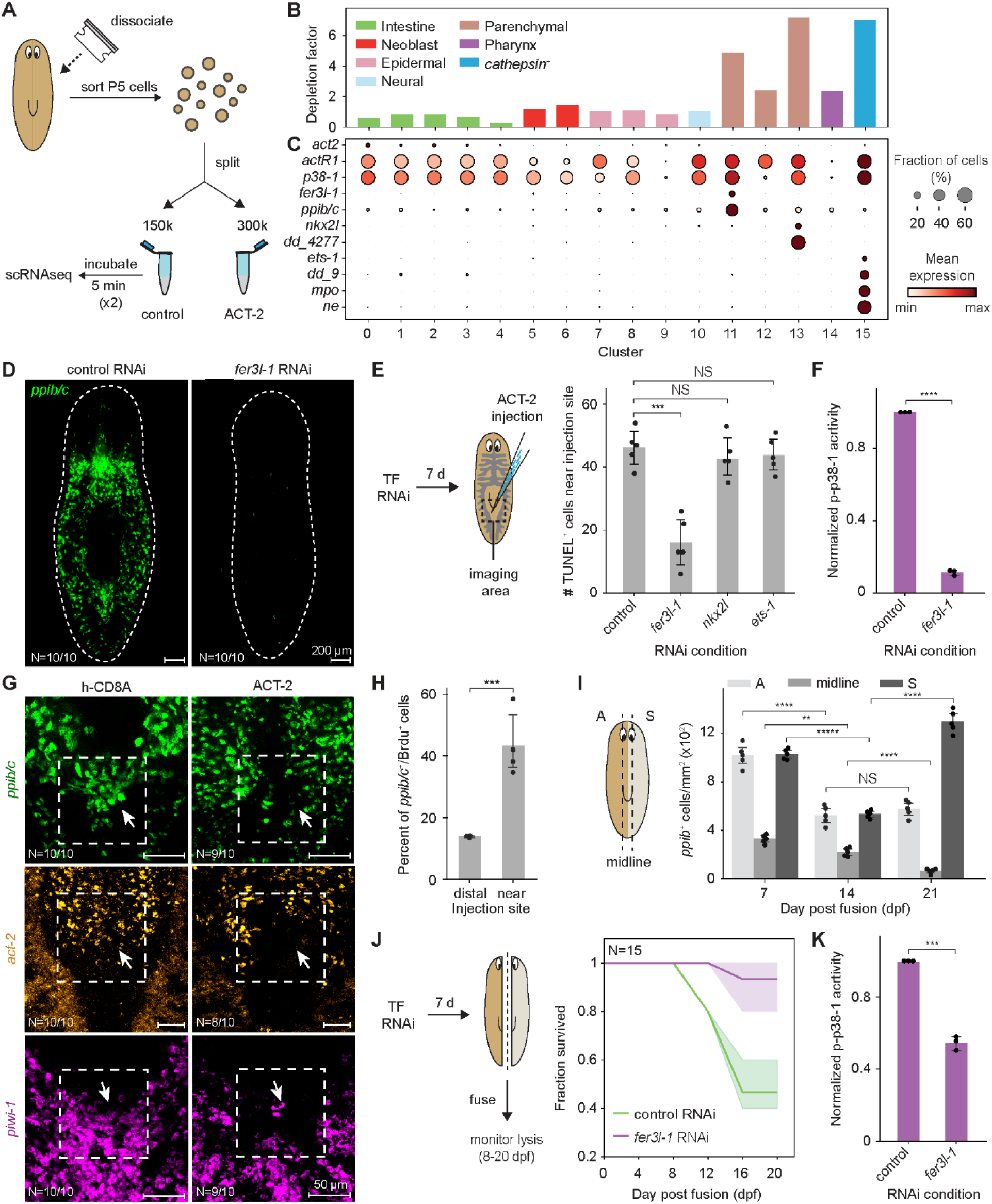
Ruptoblasts are glandular/secretory cells and mediate cytotoxicity *in vivo*. **(A)** Schematics showing the strategy to identify the molecular identity of ruptoblasts. Note the cells were incubated with h-CD8A or ACT-2 twice for 5 min each before proceeding with sequencing (see **Methods**). **(B)** Depletion factor for each cell cluster, defined as the ratio of cell abundance between h-CD8A and ACT-2 treated samples. **(C)** Dotplot showing expression of *act2*, *actR1*, and *p38-1* along with select cluster markers and immune related-genes in each cell cluster. Color: mean expression calculated across control and ACT-2 treated conditions, dot size: fraction of expressing cells. The broad expression patterns of *actR1* and *p38-1* were also consistently observed in the planarian cell atlas^23^. **(D)** FISH of the ruptoblast marker, *ppib/c* (*dd_348*), showing complete elimination of ruptoblasts after *fer3l-1* (*dd_8096*) RNAi. Dashed lines: animal outline. **(E)** TUNEL^+^ cells counted near the injection site for control, *fer3l-1, nkx2l (dd_13898),* and *ets-1 (dd_2092)* RNAi-treated animals 3 hr post ACT-2 injections. **(F)** ACT-2-induced p38-1 activation is reduced in *fer3l-1* RNAi animals at 3 hpi, with p-p38-1 levels normalized against the control. **(G)** ACT-2 injection ablate ruptoblasts (top), activin-secreting cells (middle), and neoblasts (bottom) at the injection site (dashed boxes and arrows). **(H)** Quantification of newly differentiated ruptoblasts (BrdU^+^/*ppib/c^+^*) in regions distant from and near the injection site at 5 d post ACT-2 injection. **(I)** Density of ruptoblasts in asexual (A) side, midline region where asexual and sexual cells interact, and sexual (S) side in chimeras at 7, 14, and 21 dpf. **(J-K)** *fer3l-1* RNAi reduces lesioned fraction (J) and p-p38-1 levels (K) in chimeras. N in (D, G) represents the number of animals showing the phenotype among all examined. Data points in (E, H, I) represent individual animals pooled from two independent experiments, in (F, K) represent biological replicates, with each containing three animals. SD of the survival curves in (J) is from three independent experiments with N denoting the total number of animals monitored. Statistical significance was determined by a two-sided t-test, with SD as error bars. **p < 0.01, ***p <0.001, ****p < 0.000l, NS, not significant.

To verify which cluster is responsible for ACT-2-induced cytotoxicity *in vivo*, we selectively ablated each of the three candidate populations by RNAi against their specific transcription factors (TFs) (**Figure 4D** and **S6C**) and then assessed whether these treatments suppressed ACT-2-induced cell death. Only animals treated with *fer3l-1* RNAi, which specifically targets cluster 11, exhibited a marked reduction in cell death near the ACT-2 injection site (**Figure 4E** and **S6D**), along with diminished p38 activation (**Figure 4F**). In contrast, depletion of clusters 13 and 15 via RNAi against their respective TFs, *nkx2l* and *ets-1*, had no such effect. These findings identified cluster 11 as ruptoblasts and the principal mediator of ACT-2-induced cytotoxicity *in vivo*. Notably, *cathepsin*^+^ cells (cluster 15), previously thought to be presumptive immune cells in planarians due to their bacterial engulfment capacity^37^ and expression of many immune-related genes (**Figure 4C**), such as homologs of mammalian neutrophil markers, myeloperoxidase (*mpo*)^38^ and neutrophil elastase (*ne*) – granular enzymes essential for chromatin decondensation and formation of extracellular traps^39^ – were not responsible for the observed cytotoxicity under these conditions. This result aligns with the general expectation that phagocytic and cytotoxic functions are executed by distinct cell types.

### Ruptoblasts exhibit broad cytotoxicity *in vivo*

Ruptoblasts are widely distributed throughout the planarian body, except in the head region, and are often located in proximity to activin-secreting cells and neoblasts (**Figure 4D** and **S6E)**. Injection of ACT-2 eliminated ruptoblasts at the injection site, along with nearby cells within a ∼100 μm radius, including activin-secreting cells and neoblasts (**Figure 4G** and **S6F**). This demonstrated the broad and potent killing activity of ruptoblasts *in vivo*. The lost ruptoblasts were quickly replenished by neoblasts, as shown by bromodeoxyuridine (BrdU) pulse-chase experiments (**Figure 4H** and **S6G)**. This continuous replenishment is essential for sustaining ruptoblast population and preserving their functions.

In chimeras, ruptoblast density sharply declined throughout the body (**Figure 4I** and **S6H)**, consistent with the *ex vivo* observations where both the asexual and sexual cells lysed similarly upon ACT-2 activation (**Figure S4C**). In surviving chimeras, however, ruptoblasts rebounded in sexual tissues by 21 dpf, potentially reflecting distinct steady states of the two genotypes under high activin. Importantly, eliminating ruptoblasts via *fer3l-1* RNAi before fusion prevented lysis (**Figure 4J**) and reduced p38-1 activity (**Figure 4K**) in chimeras, confirming that ruptoblasts are central to the destructive inflammatory response triggered by prolonged activin elevation.

### Ruptoblasts assist in bacterial clearance

To determine whether ruptoblasts are bona fide immune cells, we examined their roles in combating pathogens by challenging planarians with varying concentrations of pathogenic *Pseudomonas* (**Figure 5A**)^25^. We found that bacterial infection activated the activin pathway, as measured by p-Smad2/3, but only at high bacterial loads (**Figure 5B** and **S7A**). Additionally, ACT-2 enhanced bacterial engulfment by planarian cells without affecting pinocytosis (**Figure S7B**) demonstrating its specificity as a cytokine.

**Figure 5.**
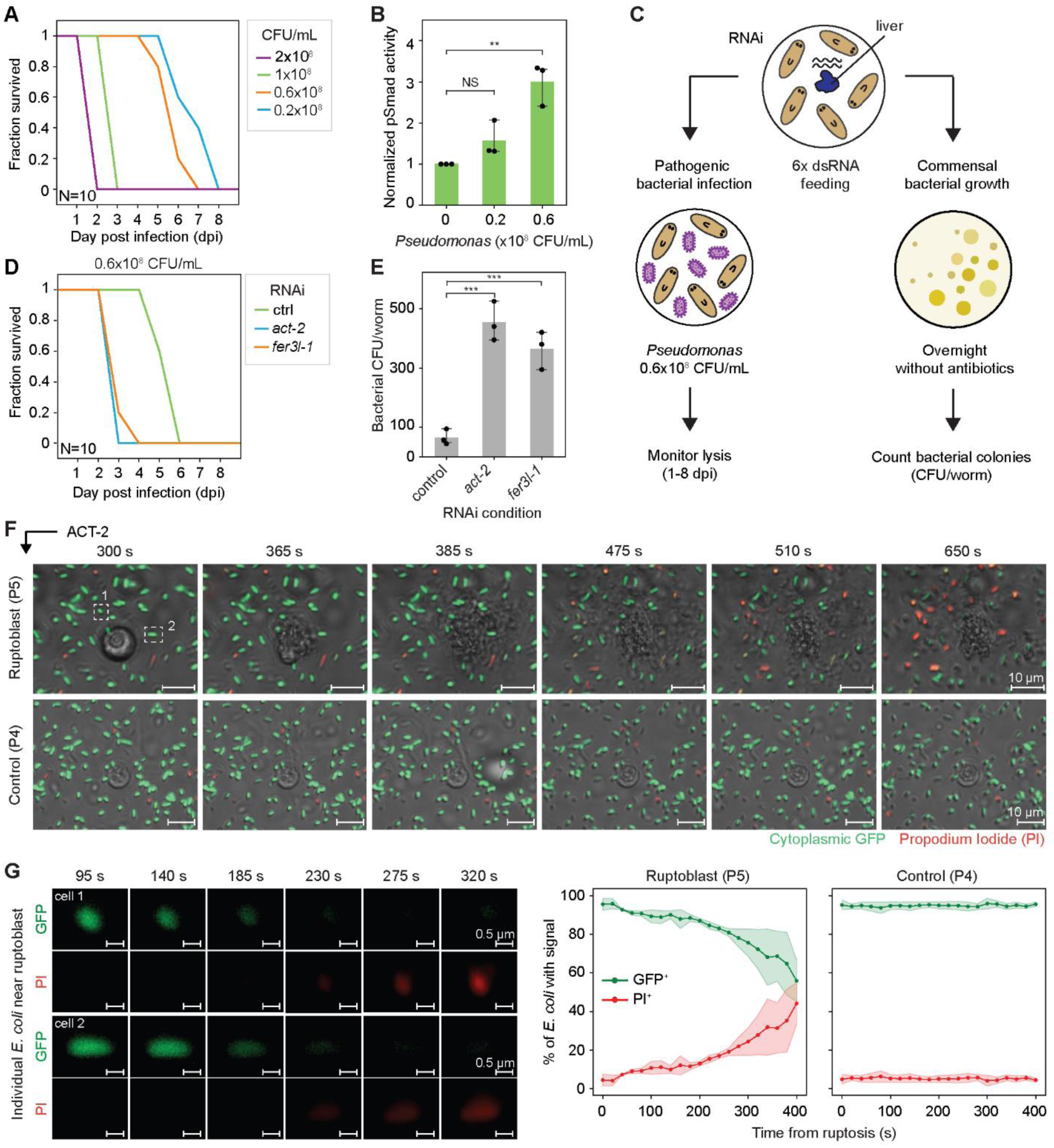
Ruptoblasts assist in bacterial clearance. **(A)** Survival curves of asexual planarians exposed to varying concentrations of pathogenic *Pseudomonas*. **(B)** p-Smad2/3 level in animals exposed to different *Pseudomonas* concentrations. **(C)** Schematics outlining the experimental workflow used to measure the function of ruptoblasts in bacterial clearance *in vivo*. **(D)** Survival curves of *act-2* and *fer3l-1* RNAi animals exposed to 0.6x10^8^ CFU/mL of *Pseudomonas*. **(E)** Quantification of commensal bacterial load in control, *act-2* and *fer3l-1* RNAi animals. **(F)** Representative snapshots showing that ruptoblasts kill nearby *E. coli* (top) upon ACT-2 whereas non-ruptoblast P4 cells (bottom) do not. *E. coli* death is evidenced by the loss of cytoplasmic GFP (green) followed by PI uptake (red). White dashed boxes highlight the two *E. coli* cells shown in F. The corresponding video is shown in **Video S4**. **(G)** (Left) Magnified snapshots of individual *E. coli* undergoing cell death following ruptosis. The loss of cytoplasmic GFP signal occurs abruptly within 45 s, followed by PI uptake in the next 1.5 min, indicating that both inner and outer membranes are permeabilized by the killing agents released by ruptoblasts. (Right) Quantification of the fractions of GFP^+^ and PI^+^ *E. coli* over time, illustrating the kinetics of bacterial death at the population level. N in (A, D) represents the total number of animals monitored. Data points in (B, E) represent biological replicates, each containing three animals. Statistical significance is determined by a two-sided t-test, with SD indicated by error bars. SD in (G) is calculated from three experiments, with time zero set at the onset of ruptosis. **p < 0.01, ***p <0.001, NS, not significant.

Both *act-2* RNAi and *fer3l-1* RNAi prior to infection increased animals’ sensitivity to bacterial challenge; these animals lysed days ahead of controls (**Figure 5C** and **5D**). The timing of lysis closely mirrored that of animals exposed to higher bacterial loads (**Figure 5A**), suggesting that infection-induced activin activates ruptoblasts to assist in bacterial clearance. Consistently, these ‘immune compromised’ animals exhibited significantly higher commensal bacterial loads compared to controls (**Figure 5C** and **5E**).

This prompted us to investigate whether ruptoblasts can directly kill bacteria. Remarkably, incubation of P5 cells with GFP-expressing *E. coli* did not induce ruptosis or bacterial death, suggesting that ruptoblasts do not sense bacteria like other immune cells^40^. However, upon adding ACT-2 to induce ruptosis, the nearby bacteria rapidly lost GFP signal, indicative of membrane leakage, and ultimately took up PI (**Figure 5F** and **5G**, **Video S4**). A single ruptoblast killed ∼45% of surrounding bacteria, and the abrupt loss of their GFP implied that the released cytotoxic agents are highly potent. Notably, we did not observe ruptoblasts engulfing bacteria, consistent with prior studies showing that planarian phagocytic cells are *cathepsin^+^* ^37^. Since *cathepsin^+^* cells also express *act-2* (**Figure S7C**), it is plausible that these phagocytes secrete activin to activate ruptoblasts, thereby enhancing the overall clearance of infected cells and pathogens. Collectively, these results indicate that activin and ruptoblast-mediated responses serve as a secondary host defense complementing phagocytic activity.

### Ruptoblasts have potent cross-species cytotoxicity

To determine whether ruptoblasts release generic cytotoxic agents or kill through a planarian-specific signaling mechanism, we examined their ability to kill mammalian cells. We co-incubated ruptoblasts with human embryonic kidney (HEK293) cells and mouse macrophages (RAW264.7). Upon activin stimulation, ruptoblasts underwent ruptosis inducing rapid death of both HEK and RAW cells (**Figure 6A**, **Video S5**), as indicated by their detachment from the surface, swelling and rounding, nuclear exposure, and PI uptake within a range of ∼200 μm from the ruptoblast. A single ruptoblast was sufficient to kill ∼60-70 target cells at a time. Additionally, ruptosis triggered pyroptosis, an inflammatory cell death mechanism^41^, in RAW cells, as evidenced by caspase-1 activation (**Figure 6B**). These results demonstrate that ruptoblast-derived cytotoxic agents are extremely potent with broad cross-species activity.

**Figure 6.**
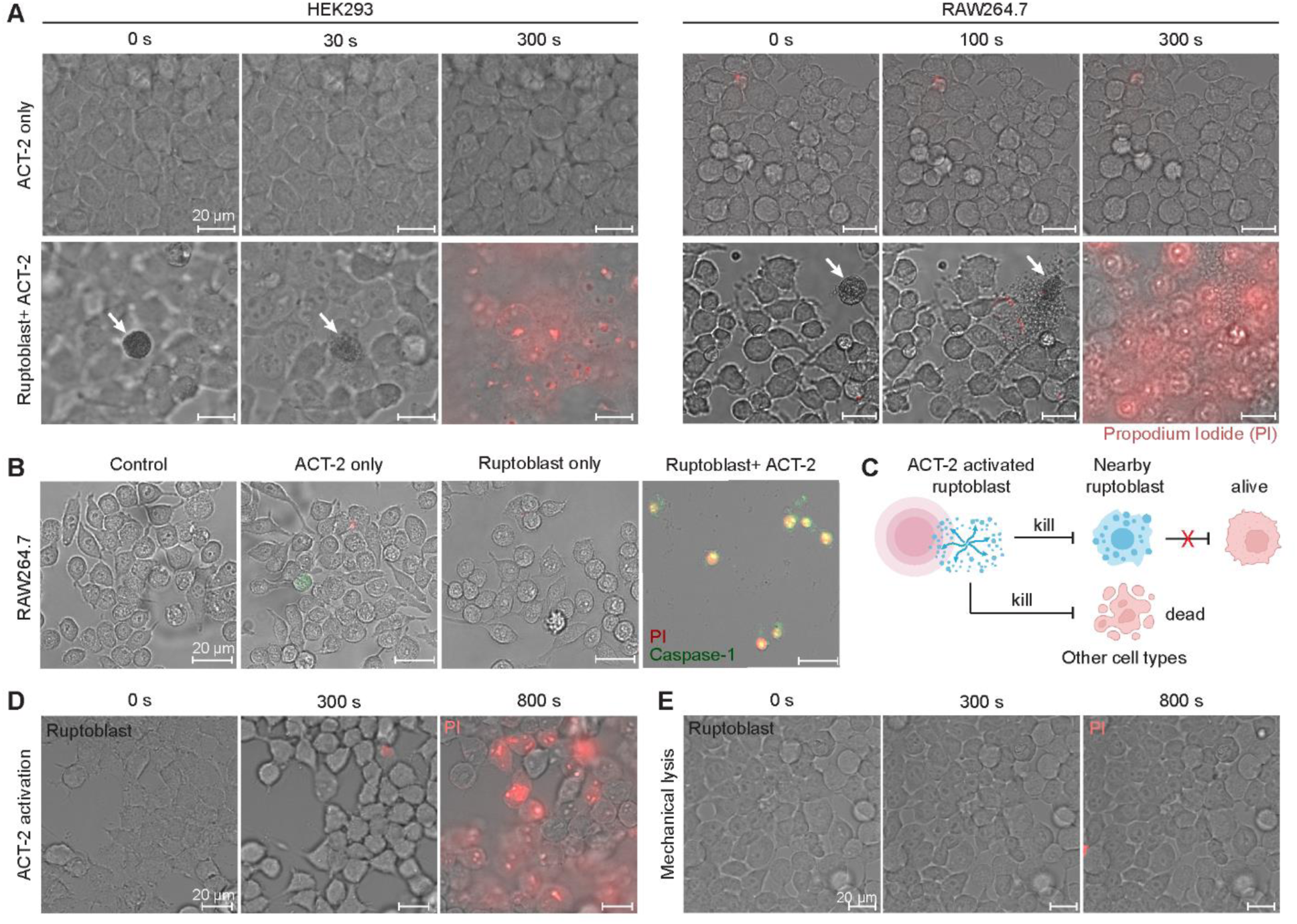
Ruptoblasts kill mammalian cells efficiently. **(A)** Snapshots of ruptoblasts (white arrows) killing HEK293 cells (left) and RAW264.7 cells (right). ACT-2 are added at time 0. The cytotoxic killing is not observed with ACT-2 added alone, confirming that the killing is mediated by the contents released from the ruptoblasts. **(B)** Ruptosis induces inflammatory pyroptosis in RAW 264.7 cells as evidenced by caspase-1 activation. This response is not observed when cells are treated with ACT-2 or ruptoblasts alone, indicating that both are required to trigger inflammatory response. **(C)** To minimize collateral damage of ruptosis, ruptoblasts killed by neighboring ruptosis must themselves be incapable of releasing cytotoxic agents, implying the activation of the killing agent requires the activin activation. **(D)** Snapshots of HEK293 cell death caused by ruptoblast-derived supernatant collected after ACT-2 induced ruptosis. The supernatant alone is sufficient to kill HEK293 cells. **(E)** Snapshots of HEK293 cells treated with supernatant from mechanically disrupted ruptoblasts without ACT-2 activation. No cytotoxic effect is observed, suggesting that the cytotoxic agents require activation by ACT-2 to convert into their active or toxic form. All corresponding videos are shown in **Video S5**.

This remarkable potency raises the question of how ruptoblast cytotoxicity is spatially and temporally contained *in vivo*, given that ruptoblasts are often located in close proximity and could kill neighboring ruptoblasts, potentially triggering a chain reaction (**Figure 6C**). To address this, we tested whether passive ruptoblast rupture is sufficient to releases active cytotoxic agents. After isolating ruptoblasts, we induced ruptosis with activin, collected the supernatant, and confirmed its cytotoxic effect on HEK cells (**Figure 6D**, **Video S5**). In contrast, supernatant from mechanically lysed ruptoblasts without activin activation did not have killing activity (**Figure 6E**, **Video S5**), demonstrating that activin-triggered modifications are necessary to generate the active form of the cytotoxic agents. Furthermore, supernatant from non-ruptoblast (P4) cells showed no detectable cytotoxicity (**Figure S7D**), and brief incubation (15 min) abolished the killing activity of ruptoblast-derived supernatant (**Figure S7E**). These findings underscore the strict spatiotemporal confinement of the ruptoblast cytotoxicity, preventing unintended propagation of cell death.

### The explosive nature of ruptosis is regulated by intracellular calcium and cytoskeletal dynamics

Ruptoblasts are distinguished by their explosive mode of action, rivaling the fastest cytotoxic mechanisms described to date. This raises a key question: what molecular machinery enables such rapid and complete cellular destruction? Given their abundant expression of calcium-binding proteins (e.g., *calretinin*, *calmodulin*, and *calcyphosine*) (**Figure S7F**) and the observation that ruptosis can occur in calcium-free media, we postulated that intracellular calcium may modulate the kinetics and magnitude of ruptosis.

Live imaging of a calcium indicator (Fluo-4 AM) revealed a sharp rise in intracellular free Ca^2+^ immediately before plasma membrane rupture (**Figure 7A** and **7B**, **Video S6**). Chelation of intracellular calcium with BAPTA-AM inhibited full membrane disruption, constraining most granules within the cell (**Figure 7C** and **7D**, **Video S6**). The few granules that were released had significantly reduced dispersal ranges (**Figure 7E**). The actin cytoskeleton also persisted, but nuclear breakdown still proceeded as in control cells. These findings indicate that intracellular calcium is not required for initiating ruptosis but essential for full cytoskeletal disassembly and plasma membrane rupture, which are key steps in converting the activin signal into explosive intracellular content release.

**Figure 7.**
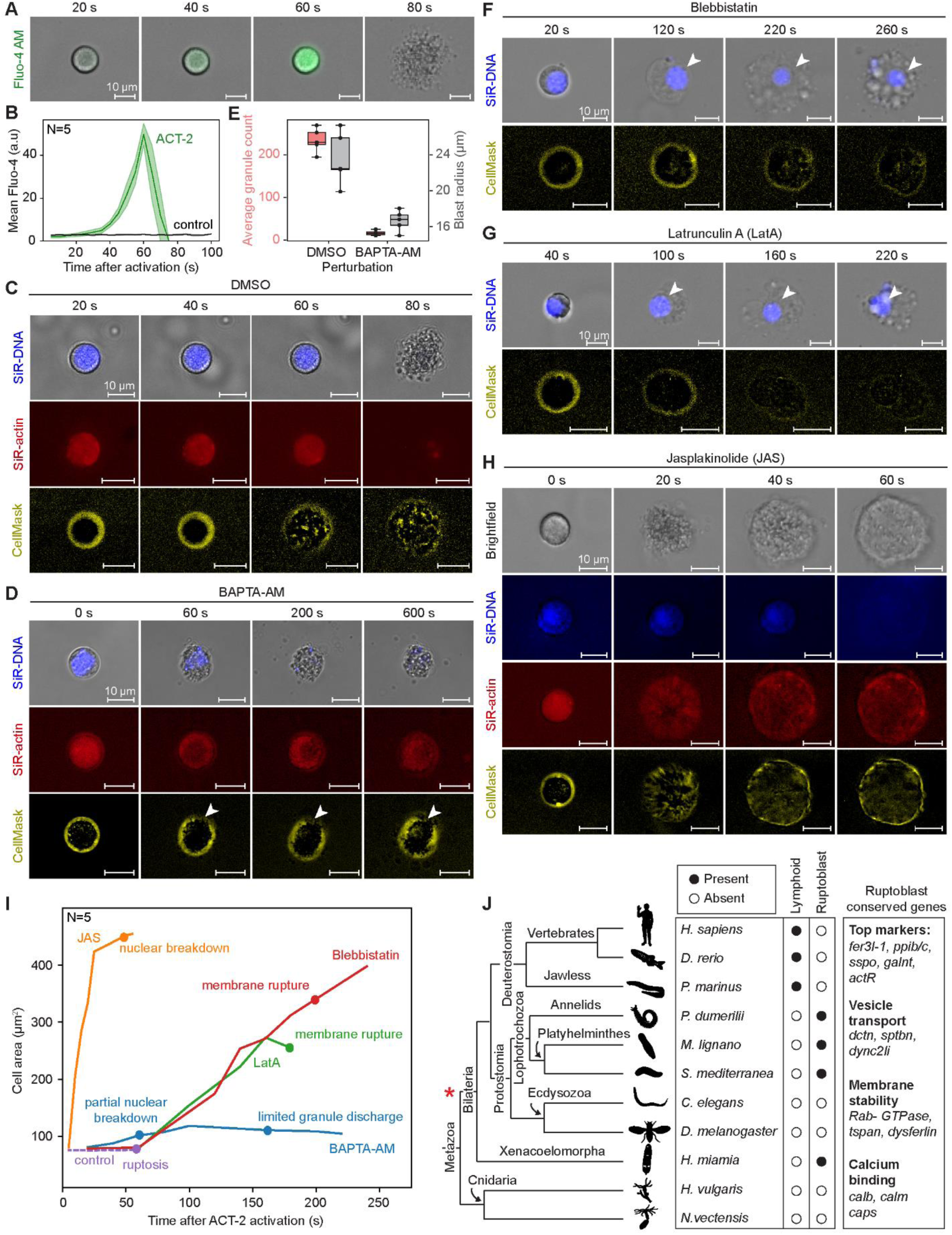
Ruptosis is regulated by intracellular calcium and cytoskeleton dynamics (A-B) Fluorescence snapshots (A) and quantification (B) showing a sharp rise in intracellular Ca^2+^ immediately preceding plasma membrane rupture following ACT-2 treatment, which is absent after control (h-CD8A) treatment. **(C)** Snapshots highlighting the breakdown of key cellular components during ruptosis in control cells. **(D)** Snapshots of ruptoblasts pretreated with the intracellular calcium chelator BAPTA-AM (in CMF). Ruptosis still initiates upon activin exposure, leading to nuclear disintegration, but only partial membrane rupture and reduced granule release. Actin cytoskeletal structure persists longer compared to controls. These changes are accompanied by slight cell expansion. White arrow: local membrane rupture. **(E)** Boxplots showing granule discharge (pink) and blast radius (grey) in control (DMSO-treated) and BAPTA-AM-treated cells. Granule count represents the average number of granules across 5 frames with the highest count per cell. Blast radius was calculated from the maximum area of granule spread observed during ruptosis. **(F-H)** Representative snapshots showing the effects of cytoskeleton perturbations on ruptosis. Treatment with blebbistatin (F) or LatA (G) led to gradual cell expansion and significantly delayed membrane rupture. The nucleus (white arrow) remains visible under both conditions. In contrast, treatment with JAS (H) leads to abrupt cell expansion. Granules are constrained, often clustering near or within the cell body, and the plasma membrane appear to reseal following granule release. **(I)** Quantification of cell area over time under different perturbation conditions, with annotations highlighting key cellular events over time. **(J)** Ruptoblast is likely an ancient cytotoxic cell type, whereas lymphoid cytotoxic cells are vertebrate specific^6–9, 48^. Filled circles: presence, empty circles: absence, based on gene expression. Red star: proposed evolutionary origin of ruptoblasts. Genes conserved in all identified ruptoblast-like cells are listed on the right. In *C. elegans* and the cnidarians *H. vulgaris* and *N. vectensis*, no *fer3l-1* homolog was found. In *H. sapiens*^49^ and the fly *D. melanogaster*^50^*, fer3l-1* is expressed predominantly in neurons, while in the zebrafish *D. rerio*^51^, the expression of *fer3l-1* is minimum. All corresponding videos for (A, C-D, F-H) are shown in **Video S6**. Fluorescence intensities were adjusted over time to visualize weak signals. N in (B, I) represents the number of independent ruptosis analyzed under each condition.

We next asked whether and how actomyosin cytoskeleton contributes to the explosiveness of ruptosis. We inhibited myosin II with blebbistatin or blocked actin polymerization with latrunculin A (LatA). Both treatments slowed ruptosis, resulting in gradual cell expansion and delayed plasma membrane rupture (at ∼200 s after adding ACT-2 with blebbistatin and ∼160 s with LatA, compared to ∼60-80 s in control cells) (**Figure 7F** and **7G**, **Video S6**). The nucleus were still visible in both conditions, indicating that actomyosin cytoskeleton is required to drive rapid and complete cellular destruction.

Conversely, stabilizing the cortical actin cytoskeleton with jasplakinolide (JAS), which prevents actin filament turnover, led to an abrupt cell expansion but suppressed granule discharge and membrane rupture (**Figure 7H** and **7I**, **Video S6**). Instead, membrane blebbing was observed. Nonetheless, nuclear disruption still occurred – albeit earlier compared to other conditions – suggesting nuclear dissolution alone is insufficient to drive full ruptosis. Together, these results indicate that ruptosis is initiated by activin, which triggers intracellular calcium signaling to drive energy buildup. The dynamic actomyosin cytoskeleton holds this energy, potentially through mechanical tension, and upon sudden disassembly, releases it explosively, rupturing the plasma membrane and leading to complete cellular destruction.

## Discussion

In this study, we discovered and characterized a new type of cytotoxic cell, ruptoblasts, and described a previously unknown form of cell death, ruptosis. Ruptoblasts respond specifically to activin, which functions both as a hormone and a cytokine in planarians, undergoing ruptosis and releasing diffusible cytotoxic agents capable of eliminating any nearby cells. Because both the sensing and killing functions of ruptoblasts rely on diffusible cues, the effective killing range should inherently match the activin-sensing distance. Importantly, the passive death of ruptoblasts does not release cytotoxic agents, thus avoiding chain reactions between ruptoblasts and minimizing collateral damage. Unlike conventional immune cells, which are activated by pathogens or other immune insults, ruptoblasts are triggered by activin, highlighting a regulatory mechanism finely attuned to hormonal cues, which parallels the hormonal surveillance function of mammalian cytotoxic T cells^10, 11^.

As a pro-inflammatory cell type, ruptoblasts may engage a broader immune network. They are likely activated by upstream signals from cells that recognize non-self genotypes and by *cathepsin^+^* phagocytes that detect and engulf bacteria^37^, both may producing activin (**Figure S7G**). Ruptoblasts should provide regulatory feedback to phagocytes for enhanced pathogen clearance, as evidenced by compromised infection resistance following ruptoblast ablation, whereas blocking ruptosis alone by *p38-1* RNAi only reduces inflammation^25^. In addition, analogous to immune exhaustion or other regulatory adaptations of mammalian T cells^42^, ruptoblasts may develop tolerance to sustained, yet moderate activin levels. Surviving chimeras, for example, had smaller ruptoblasts at lower densities (**Figure S6H**), indicating a shift toward a less active or immature state. These nuanced regulations invite deeper investigation into the molecular and cellular mechanisms regulating ruptoblast behavior.

Are ruptoblasts unique to planarians? Our sequencing data indicate that they lack canonical immune markers, except for *ppib/c*, a cyclophilin-type peptidyl-prolyl cis-trans isomerase known to activate T and B cells in mammals^43^, and do not express known cytotoxic effectors. This is consistent with the fact that ruptoblasts belong to a glandular/secretory cell type family^23, 24^ distinct from classical immune cells and that they execute cytotoxic killing through unique mechanisms. Strikingly, homologs of *fer3l-1* and another ∼30 highly expressed ruptoblast genes, including activin receptor, membrane stability proteins, calcium-binding proteins, and cell-type markers such as *ppib/c*, *sspo*, and *galnt* (**Figure S7F**), co-express in specific cell types across other flatworms^44^, annelids^45^, and acoel worms that are sister to all bilaterians^46^, yet appear absent in cnidarians (e.g., *Hydra* and *Nematostella*) and well-studied models (e.g., vertebrates, flies, and nematodes). This implies that ruptoblasts may represent an ancient, bilaterian-specific immune cell type subsequently lost in ecdysozoans and deuterostomes (**Figure 7J**). Their unique killing mechanism may address the special demand of regenerative animals, prevalent among Lophotrochozoa, enabling simultaneous elimination of both hormone-secreting cells and their stem cell precursors. These animals are also capable of recovering from the damages caused by ruptosis through regeneration. Investigating the functions and regulation of ruptoblast-like cells in other species is a compelling avenue for future research.

Altogether, our findings highlight the immense diversity of immune innovations across the animal kingdom and emphasize the need for broader exploration of immune systems beyond conventional models. Deeper understanding of the molecular details of ruptosis will not only reveal new cell biology paradigms but also inspire strategies for engineering cytotoxicity for enhanced therapeutics against pathogenic bacteria and malfunctioning cells.

## Limitations of study

First, understanding precisely how activin-p38 signaling, cytoskeletal components, and calcium-modulated biochemical pathways are orchestrated to produce the explosive rupture during ruptosis may open up a new area in cell biology and likely requires novel theoretical frameworks and direct mechanical measurements to fully elucidate this extremely rapid cellular process. In addition, comparing ruptosis with NETosis, another form of sacrificial cell death^33, 34, 47^, can reveal how cellular machinery yield distinct cell-death dynamics. Second, despite the remarkable potency and broad-spectrum efficacy of ruptoblast-derived cytotoxic agents, the precise identity of the cytotoxic agent remains unknown. Pilot experiments narrowed the active agent to a protein of a modest size (30-100 kDa) (**Figure S7H** and **S7I**), with activity dependent on activin-induced modifications. Although this information provides an entry way, identifying this specifically modified, highly unstable protein presents a technical challenge. Nonetheless, elucidating the molecular switch underlying this activation-dependent cytotoxicity represents a promising direction for harnessing its killing activity for future applications.

## Supplemental Information

**Figure S1.**
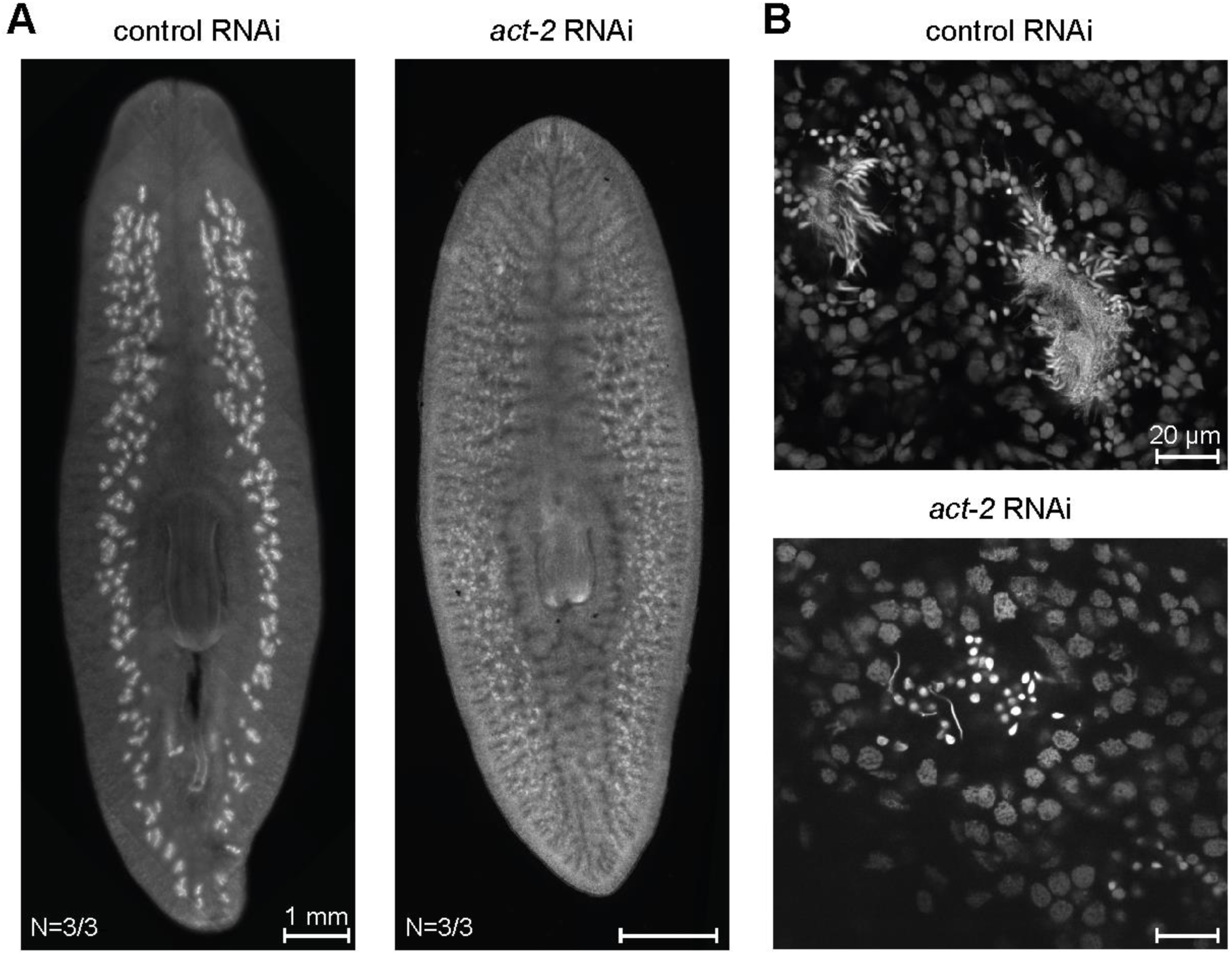
*act-2* is required for spermatogenesis in sexual planarians. **(A)** DAPI-stained testes of sexually mature planarians following 6 feedings of control or *act-2* RNAi. In control RNAi animals, large testes are distributed beneath the dorsal epithelium, whereas *act-2* RNAi animals exhibit only small clusters. N denotes the number of animals displaying the reported phenotypes out of total analyzed. **(B)** Higher magnification imaging of testis lobules in control and *act-2* RNAi reveals defects in spermatogenesis. In controls, individual testis can be identified by the accumulation of sperm in the center of the lobule with a long, string-like morphology, surrounded by cells at various stages of spermatogenesis. In contrast, testis lobules in *act-2* RNAi animals are smaller, largely devoid of differentiated cells across spermatogenic stages.

**Figure S2.**
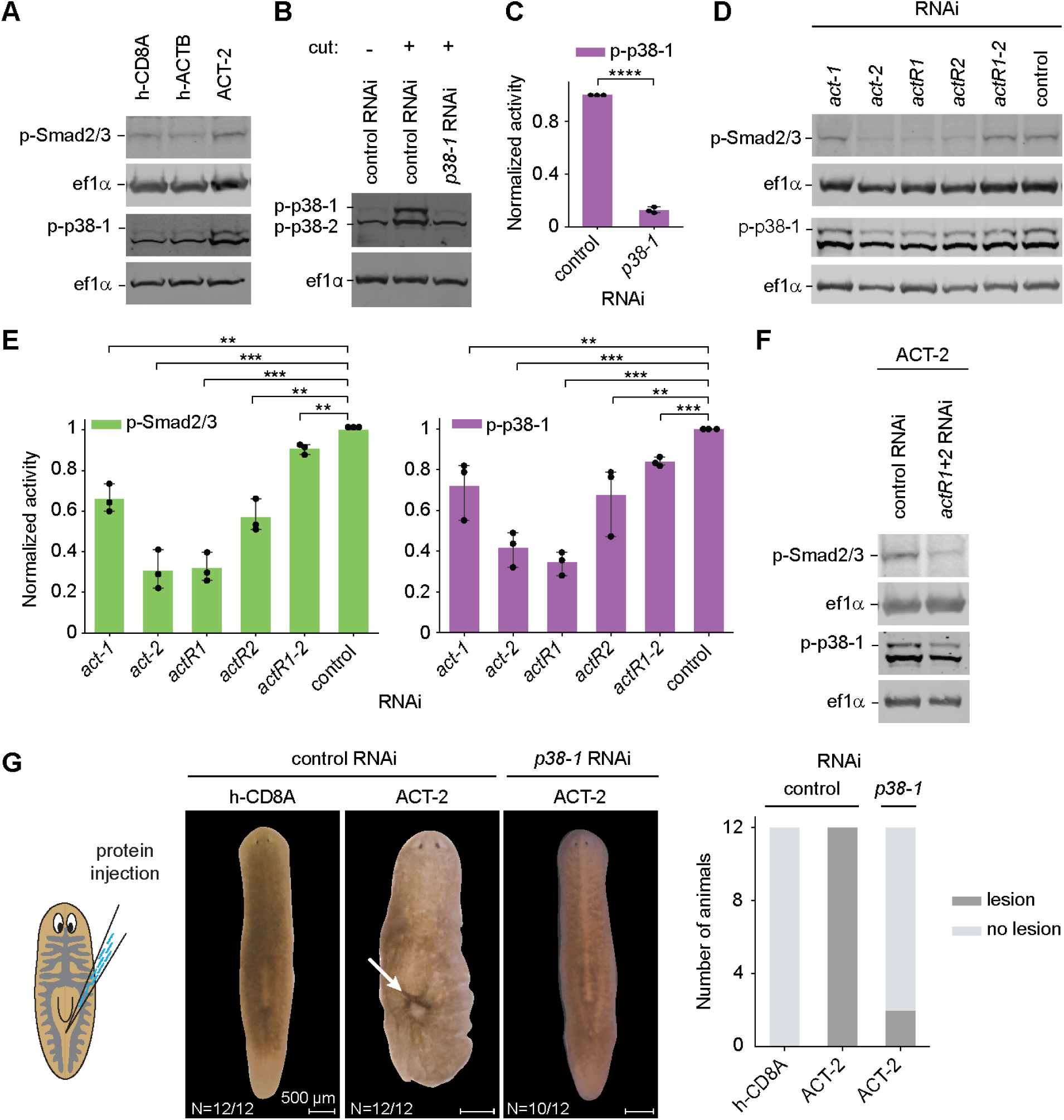
p38-1 is downstream of activin. **(A)** Representative Western blotting images showing the increase of p-Smad2/3 and p-p38-1 in animals injected with ACT-2 proteins, but not with control proteins (h-CD8A, h-ACTB). Quantification is shown in Figure 1C. **(B-C)** *p38-1* RNAi eliminates the p-p38-1 band after amputation, validating the specificity of the antibody. Note that amputation is used to activate p-38 as described^25^. **(D-E)** Western blotting images (D) and quantification (E) showing that RNAi of the two activin homologs and three activin receptor homologs significantly reduce p-Smad2/3 (green) and p-p38-1 (magenta) levels. Quantified fold activation normalized to the levels in control RNAi animals. Statistical significance was determined by a two-sided t-test, error is reported as SD from three independent experiments, each containing five animals. **p < 0.01, ***p <0.001, NS, no significant difference. **(F)** Representative Western blotting images showing knockdown of *actR1* and *actR2* blocks the effect of ACT-2 injection. Quantification is shown in Figure 1D. **(G)** (Left) Brightfield images of control RNAi animals injected with either control (h-CD8A) or ACT-2 protein, and *p38-1* RNAi-treated animals injected with ACT-2 protein. White arrow indicates lesions formed following injection. (Right) Number of animals with and without lesions across conditions.

**Figure S3.**
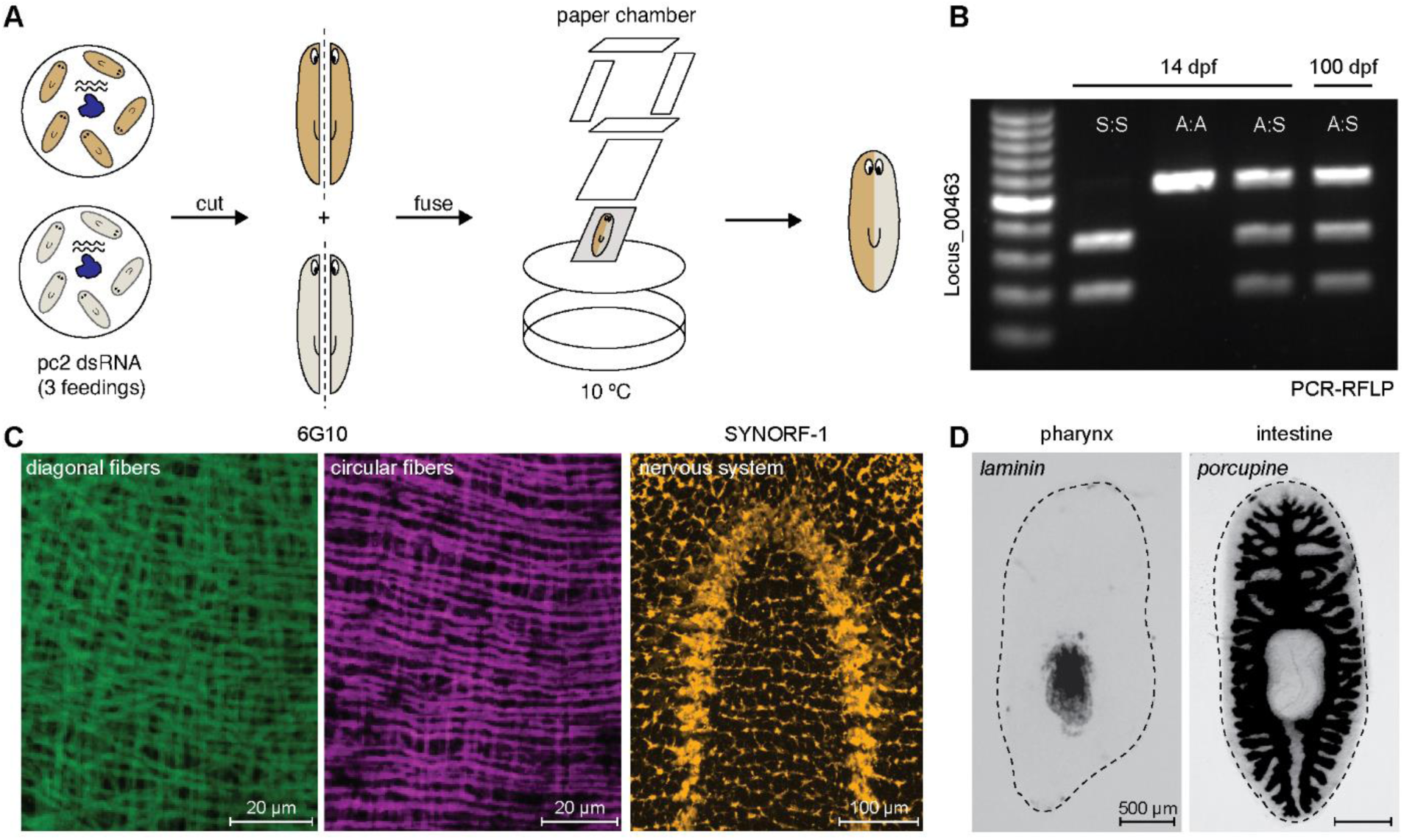
Complete tissue integration in genetic chimeras. **(A)** Schematics showing the workflow of fusion experiments. Asexual and sexual animals are fed dsRNA against *pc2*, and the opposing halves are aligned and covered with Zig-Zag cigarette rolling paper. The fusion was further constrained by four pieces of filter paper, and a Kimwipe soaked with IO water to prevent drying. **(B)** PCR-RFLP analysis of chimeras at 14 dpf and 100 dpf. Genotyping is based on locus 00463 cut by ScaI in sexual cells but not in asexual cells. **(C)** Immunofluorescence showing complete structural integration of muscle fibers and neural tissue at the midline in chimeras at 20 dpf. **(D)** WISH images of *laminin* (left) and *porcupine* (right) at 20 dpf illustrating the integration of pharynx and primary gut branches across the midline.

**Figure S4.**
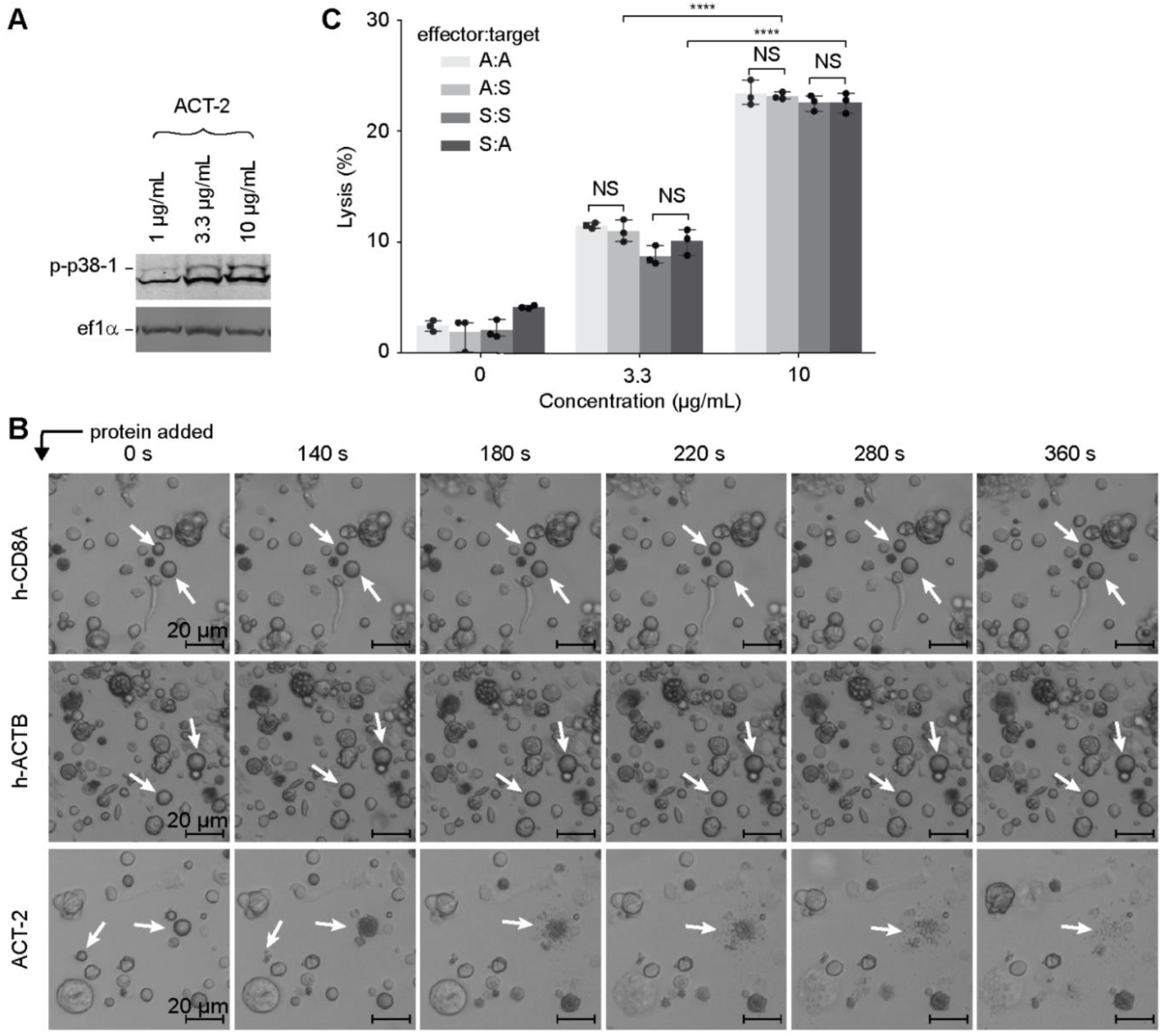
ACT-2 induces cell lysis regardless of the genotype. **(A)** Representative Western blotting image showing that p-p38-1 level increases after activin treatment in a dose-dependent manner. Quantification is shown in Figure 3B. **(B)** Snapshots showing cells undergoing explosive lysis only when incubated with 10 μg/mL ACT-2, but not with control proteins (h-CD8A or h-ACTB). White arrows indicate potential ruptoblasts based on morphology. The corresponding video is shown in **Video S1**. **(C)** Flow cytometry analysis measuring PI^+^ cells to assess lysis in response to distinct genotypes alone or in the presence of 3.3 or 10 µg/mL of ACT-2. Asexual and sexual cells were labeled with CFSE and CellTrace Far Red, respectively, mixed at a 1:1 ratio (200,000 cells total), and analyzed across all labeling combinations. ACT-2 alone is sufficient to induce lysis. Error bars: SD from three independent experiments. ***p <0.001, ****p <0.0001, NS, no significant difference, two-sided t-test.

**Figure S5.**
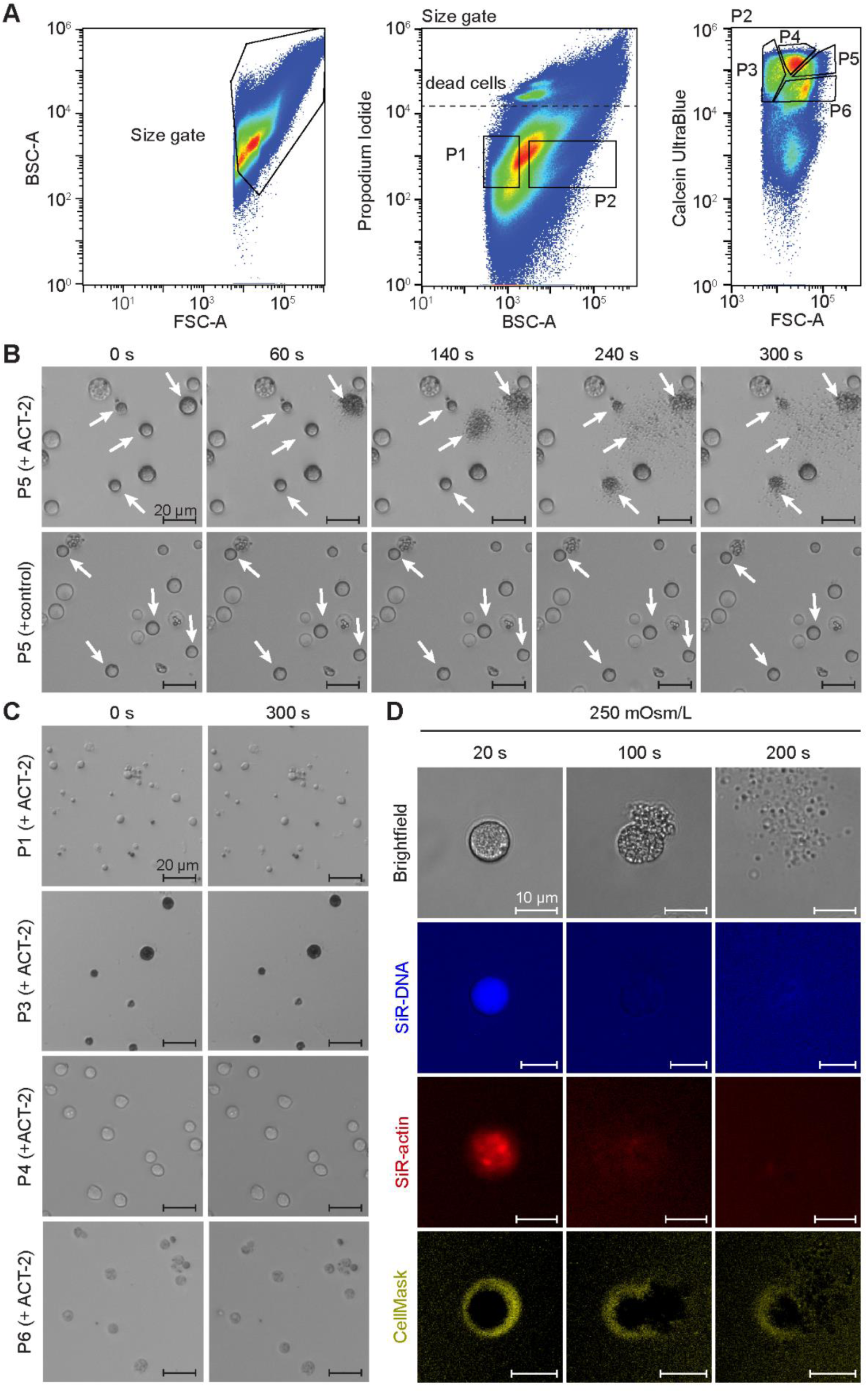
Sorting strategy to enrich for ruptoblasts. **(A)** Flow cytometry gating strategy for sorting ruptoblasts. (Left) Initial gating based on size (Forward scatter: FSC) is used to eliminate debris and contaminants. (Middle) A second refinement step uses granularity (Back scatter: BSC) and PI stain to exclude low-granularity cells (P1) and dead cells. (Right) P2 population is further subdivided into distinct subpopulations (P3, P4, P5, P6) based on Calcein UltraBlue intensity and FSC. **(B-C)** Bright field imaging snapshots showing sorted P5 (B) and all other subpopulations (C) after exposure to 10 µg/mL ACT-2 protein. P5 is the only population that responds to ACT-2. In addition, P5 population incubated with control protein (h-CD8A) do not undergo lysis. White arrows indicate potential ruptoblasts based on morphology. The corresponding video is shown in **Video S1**. **(D)** Snapshots showing ruptosis proceed normally in CMF media with sucrose at 250 mOsm/L.

**Figure S6.**
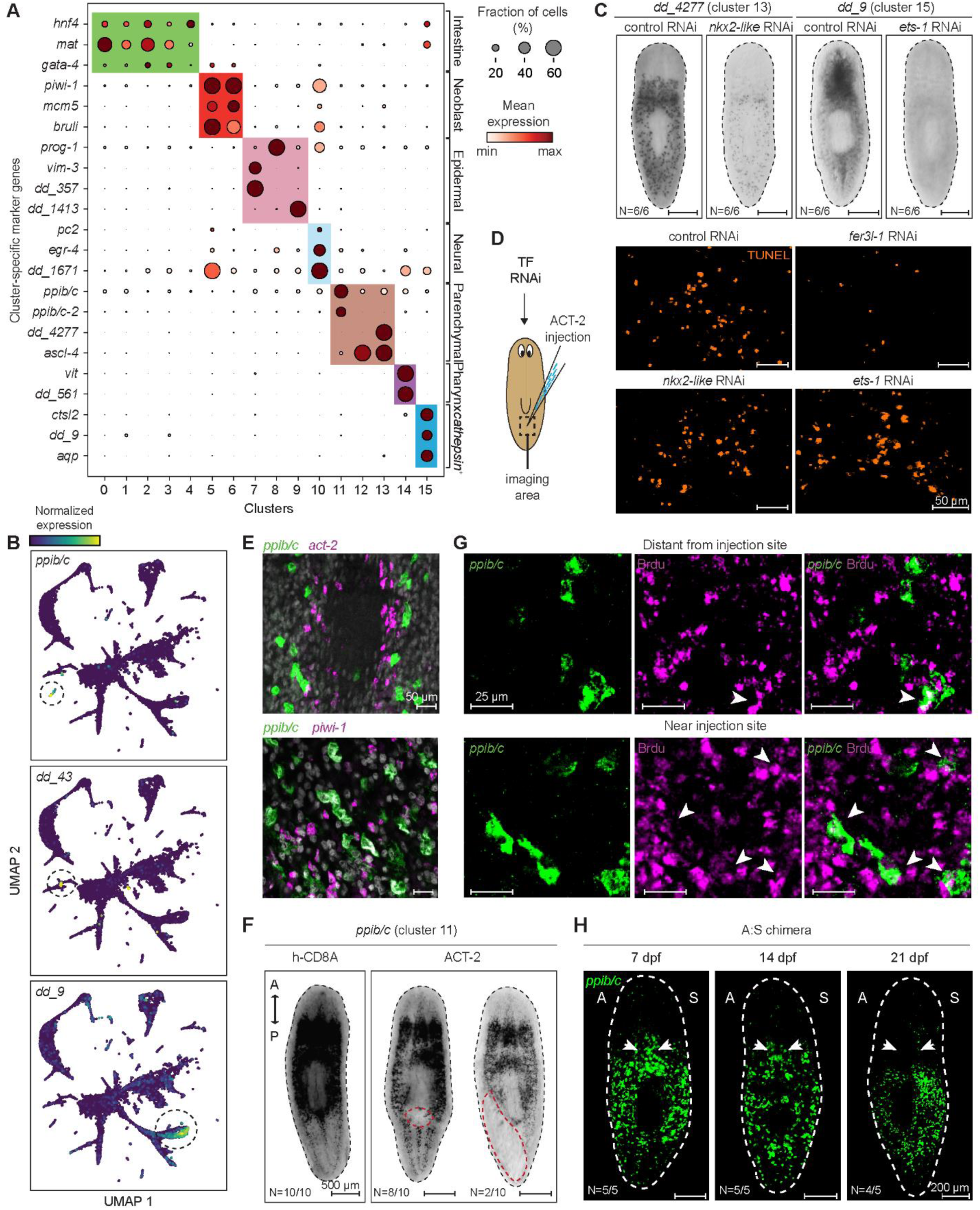
Molecular identity of ruptoblasts. **(A)** Dotplot showing expression of selected marker genes used in annotating cell type of the clusters. Contig numbers for all mentioned genes are listed in **Table S2**. **(B)** Marker gene expression of cluster 11, 13 and 15 overlaid on the UMAP projection of planarian cell atlas^23^. Dashed circles indicate cells with high marker expression, demonstrating the specificity of these markers in the full atlas. **(C)** WISH images showing the expression of *dd_4277* in control and *nkx2l* (*dd_13898*) RNAi (left) and *dd_9* in control and *ets-1* (*dd_2092*) RNAi (right) animals suggesting that knockdown of these TFs eliminate the corresponding cell populations. Dashed lines: animal outline. **(D)** Representative images showing TUNEL^+^ cells near the ACT-2 injection site, which is quantified in Figure 4E, in control, *fer3l-1, nkx2l* and *ets-1* RNAi animals. Dashed box: imaging area. **(E)** Double FISH showing that ruptoblasts, activin-secreting cells (*act-2^+^*) and neoblast (*piwi-1^+^*) are in close proximity. **(F)** WISH images showing the elimination of *ppib/c^+^* cells (cluster 11) upon 10 μg/mL ACT-2 protein injection into the parenchymal tissue between posterior gut branches. Red dashed lines denote the region where cells have been removed, and black dashed lines indicate animal surface. **(G)** Representative images of BrdU labeling (magenta) to detect newly differentiated ruptoblasts (BrdU^+^/ *ppib/c*^+^) at 5 dpi in regions distant from and near the injection sites. Corresponding quantification plot is shown in Figure 4H. **(H)** FISH showing the distribution of ruptoblasts (*ppib/c*) in chimeras at 7, 14 and 21 dpf. A: asexual side, S: sexual side, dashed lines: animal outline. White arrows highlight the decrease in ruptoblast density along the midline. In (C), (F), and (H) N denotes the number of animals consistent with the images out of the total number of animals analyzed.

**Figure S7.**
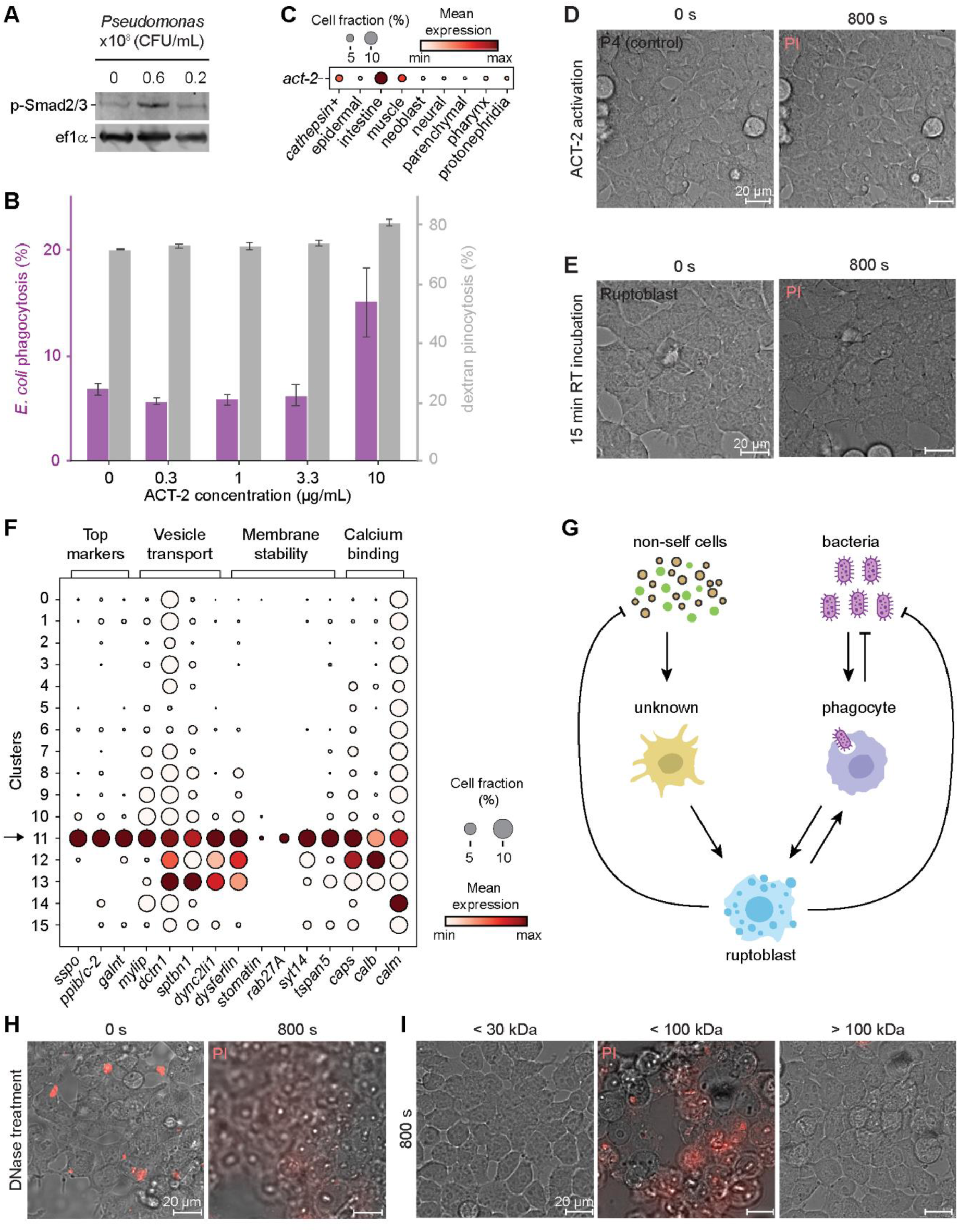
Ruptoblasts operate within a broad immune network. **(A)** Representative Western blotting image showing *Pseudomonas* infection activates the activin pathway. Quantification is shown in Figure 5B. **(B)** Quantification of phagocytosis and pinocytosis of total cells in response to ACT-2. Phagocytosis of pHrodo Green *E. coli* bioparticles (magenta) and pinocytosis of dextran (grey) were measured following treatment with varying concentrations of ACT-2 protein. Error bars represent standard deviation (SD) from three independent experiments, each containing 50,000 cells. **(C)** Dotplot showing expression of act-2 in major cell types in the planarian cell atlas^23^. **(D)** Snapshots of HEK293 cells treated with supernatant from P4 cells (control) after ACT-2 exposure. No cytotoxic effect is observed, confirming that only ruptoblasts can release cytotoxic agents. **(E)** Snapshots of HEK293 cells treated with supernatant of ruptoblasts incubated at room temperature (RT) for 15 min. The rapid loss of cytotoxic activity highlights the short lifetime of the cytotoxic agent released by ruptoblasts. **(F)** Dotplot showing expression of top marker genes, vesicle transport motor proteins, regulators of membrane stability, docking, exocytosis and calcium binding proteins in ruptoblasts (cluster 11), indicated by the arrow. Contig numbers for all mentioned genes are listed in **Table S2**. **(G)** Schematics showing the immune network in which ruptoblasts are activated by cells that detect non-self cells or by phagocytes that engulf bacteria, triggering a pro-inflammatory cascade with ruptoblasts eliminating non-self cells or bacteria. Ruptoblasts also provide feedback to phagocytes, potentially enhancing their activity to clear bacteria and debris. **(H-I)** Snapshot images of HEK293 cells treated with ruptoblast supernatant following DNase treatment (H), or molecular weight cutoff filtration (I). DNase treatment did not affect cytotoxicity, excluding DNA as the killing agent. Size-based filtration using MWCO ultra centrifugal filters revealed that the cytotoxic activity is retained within the 30-100 kDa range, consistent with proteins of modest molecular weight.

## Acknowledgements

We thank Hanh Vu and Jochen Rink for sharing the ef1α antibody and stimulating discussions, Christopher Arnold for sharing the *Pseudomonas* bacteria, Uri Alon for critical discussions, Jesse Gibson, Livia Wyss, Margarita Khariton, Christopher He, Kelli Ann Lynch, Abby Thurm, Allen Yesin and Michelle Tai for experimental help and all Wang group members for feedback. C.C. is supported by a NSF Graduate Research Fellowship, a Stanford Graduate Fellowship, and a Stanford DARE fellowship. This work is supported by an NIH grant (1R35GM138061) to B.W. and a HFSP grant (RGY0085/2019) to B.W. and B.R. Some schematics were created with BioRender.com.

## Author contributions

CC, BR, and BW designed the research. CC, ES, and LZ performed *ex vivo* experiments. CC performed sequencing experiments and analyzed the data. CC, SRS, and DNS performed functional experiments. OY helped recombinant ACT-2 protein production. CJW provided advice on bacterial cell killing assay. HRT provided advice on cell biology experiments. CC and BW wrote the paper with input from all other authors. BR and BW acquired funding and supervised the project.

## Declaration of interests

The authors declare no competing interests.

## STAR METHODS

### Resource availability

#### Lead contact

Further requests for all resources and reagents in this study should be directed to and will be fulfilled by the lead contact, Bo Wang.

#### Materials availability

Generated ACT-2 recombinant protein will be made available upon request.

#### Data Availability

Raw and processed bulk RNA and scRNA-seq datasets generated in this study are available from NCBI BioProject with accession number PRJNA1233892 (bulk RNAseq) and PRJNA1233887 (scRNAseq). Any additional data required to reproduce or re-analyze will be made available from the lead authors upon request.

### Experimental model and study participant details

#### Animal maintenance

Asexual (CIW4) and sexual (S2F8b) *S. mediterranea* were maintained in the dark at 18 °C in 0.5 g/L Instant Ocean Sea Salts (IO) supplemented with 0.1 g/L sodium bicarbonate (asexual planarian media) and 0.75× Montjuic salts (sexual planarian media), respectively. Planarians were fed calf liver paste weekly, and starved for 7 days prior to all experiments.

### Method details

#### ACT-2 recombinant protein

Between human activin (h-ACTB) and the planarian ACT-2, sequence conservation is restricted to the TGF-*β* domain with only 29% amino acid identity within the domain and no homology detected elsewhere. Given this low similarity, h-ACTB is unlikely to functionally substitute for ACT-2. Therefore, we produced ACT-2 recombinant protein using a yeast expression system (Cusabio, Cat#CSB-EP3606GOQ1) to ensure proper glycosylation of the secreted protein. A 6× His-tag was added to the N-terminus. The protein was assessed by SDS-PAGE to have 88% purity. Endotoxin level, measured using the Limulus amebocyte lysate (LAL) assay, was below 1.0 endotoxin unit (EU) per microgram of protein.

#### Protein injection

20 μg/mL of control proteins – h-CD8A (Cusabio, Cat#YP004966HU) or h-ACTB (SinoBiological, Cat#10429-HNAH) – or ACT-2 protein (20 μg/mL) was loaded into needles pulled from glass capillaries (WPI, Cat#1B00F-3) on a Sutter P97 needle puller with the following settings: pressure = 500, heat = 758, pull = 50, velocity = 70, time = 200. Needles were mounted on a Sutter XenoWorks injection system, and tips were carefully opened with forceps. Animals were positioned ventral-side up on a moist filter paper placed on a cooled block and injected along the tail midline until a visible bolus formed and ceased expanding. Following injection, animals were allowed to recover in the dark at 18 °C before downstream processing.

#### Western blotting

Animals were fixed in 100 mM ZnCl_2_ fixative for 1 hr at 4 °C^52^. Lysate was obtained by homogenizing ∼3-5 animals in urea-based lysis buffer (9 M urea, 2% SDS, 130 mM DTT, 10× protease inhibitor cocktail, 1× phosphatase, benzonase) using an electronic pestle homogenizer, and incubated for 30 min, and spun down at ∼ 22,000 × g at room temperature (RT) for 10 min. Protein concentrations were determined by absorbance at 280 nm on a Nanodrop.

Samples were mixed with 1× lithium dodecyl sulfate (LDS) buffer and denatured at 70 °C. SDS-PAGE was performed using NuPAGE Novex 4-12 % gels, and proteins were transferred to nitrocellulose membranes via either wet transfer blot module (Invitrogen) or iBlot2 dry transfer (Invitrogen). The blot was blocked in Intercept blocking buffer (LI-COR), then incubated overnight with primary antibody (1:1,000 anti-p-Smad2, Cell Signaling Technology; 1:100,000 anti-ef1α, a gift from Jochen Rink’s lab generated at MPI-CBG; 1:1000 anti-p-p38, Cell Signaling Technology) in Intercept T20 (TBS) antibody diluent solution (LICORbio, Cat# 927-65001). After washing, membranes were incubated with secondary antibodies (1:20,000 anti-rabbit IRDye 800CW, LI-COR; 1:10,000 anti-mouse-Igg1-555, Biotium) in Intercept T20 (PBS) antibody diluent solution (LICORbio, Cat# 927-75001) for 1 hr. Final washes were in PBS supplemented with 0.1% Tween-20 followed by PBS alone.

Blots were imaged with a LI-COR Odyssey imager at 800 and 680 nm, and Typhoon 9500 at 555 nm. Images were quantified using ImageJ gel plugins. p-Smad2/3 and p-p38-1 levels were normalized across samples to ef1α, and fold activation of pSmad2/3 and p-p38-1 were determined by normalizing to their respective control conditions.

For performing Western blotting on cells, the same procedure was followed, except that cells were directly lysed in the urea-based lysis buffer. A 1:50,000 dilution of anti-ef1α and 1:5,000 dilution of anti-mouse-Igg1-555 were used.

#### Planarian fusion

Sexual and asexual planarians (∼3 mm in length) were fed double stranded RNA (dsRNA) against *pc2* every 4 d for 3 feedings to immobilize animals^53, 54^ required for the fusion procedure. With this body size, sexual planarians do not develop gonads. Starved animals were cut longitudinally along the midline on an iced Coolrack with weigh paper. To fluorescently label tissues, animals were soaked overnight in CellTrace CFSE (1:1,000, Invitrogen, Cat# C34554) or Celltrace Far Red (1:1,000, Invitrogen, Cat# C34564). Opposite halves of the asexual and sexual worms were pushed together and covered with Zig-Zag cigarette rolling paper, and constrained by four pieces of filter paper and a Kimwipe soaked with asexual planarian media on the side (**Figure S3A**). Fusions were left overnight at 10 °C and transferred to 1:1 mixture of Montjuic and IO salts with 1:1,000 gentamicin.

#### Genotyping

Genomic DNA was extracted as previously described^55^. Restriction Fragment Length Polymorphism (RFLP-00463) loci^30^ were amplified from genomic DNA using Phusion Polymerase with 35-40 PCR cycles. PCR products were purified, digested with ScaI-HF restriction enzyme for 2 hr, and ran on a 1% agarose gel.

#### Staining

Riboprobes for in situ hybridization were synthesized as previously described^53^. Gene fragments were amplified from cDNA using oligonucleotide primers listed in **Table S2**, and cloned into pJC53.2 (Addgene Plasmid ID: 26536).

RNA WISH and FISH were performed as described^56^. Briefly, planarians were relaxed on ice, killed in 5% N-Acetyl Cysteine (NAC) for 5 min, then fixed for 2 hr in 4% formaldehyde supplemented with 1% NP-40 at RT. After dehydration in methanol (stored at -20 °C if needed), planarians were rehydrated and bleached for 2 hr in the bleaching solution (5% formamide, 0.5× SSC, 1.2% H_2_O_2_) under bright light. They were permeabilized with 2 μg/mL proteinase K for 10 min, and post-fixed with 4% formaldehyde. Hybridization was performed at 56 °C overnight. Detection used either NBT/BCIP for WISH or tyramide signal amplification (TSA) for FISH.

For immunostaining, planarians were relaxed on ice, killed in 2% HCl for 5 min, fixed for 6 hr at 4 °C in 4% formaldehyde, and bleached overnight in 6% H_2_O_2_. Samples were blocked in 1% (w/v) BSA in PBSTx for 4 hr, incubated overnight with primary antibodies (1:1,000 6G10, DSHB; 1:1,000 anti-SYNORF1, 3C11, DSHB), followed by 1 hr blocking. Secondary antibodies (1:1,000 goat-anti-mouse IgG+IgM conjugated with peroxidase) were added overnight, and detected by TSA.

For TUNEL staining^57^, planarians were relaxed on ice, killed in 5% NAC for 10 min, fixed in 4% formaldehyde for 20 min at RT, and bleached in 6% H_2_O_2_ overnight under bright light. They were incubated in TdT enzyme mix for 4 hr at 37 °C, followed by stop/wash buffer for 10 min, and rhodamine-labeled antibody for 4 hr at RT. TdT enzyme mix, stop/wash buffer, and rhodamine antibody were obtained from Millipore ApopTag Red Kit (Cat# S7165).

BrdU labeling with RNA-FISH used a protocol described in Ref. 27. Briefly, 2 μL of BrdU stock (20 mg/mL in 50% DMSO, Roche, Cat# 10280879001) was mixed with 8 μL of calf liver paste and fed to planarians. During the BrdU chase, animals were maintained in 5× IO salts. They were then killed in 5% NAC, fixed in 4% formaldehyde for 30 min at RT, bleached in 6% H_2_O_2_ in methanol. After standard RNA FISH, samples were treated with 2N HCl containing 0.3% Triton X-100 for 30 min at RT, washed in PBSTx, and blocked in 5% RWBR and 5% horse serum for 2 hr at RT. Samples were incubated with anti-BrdU (Abcam ab6326, 1:1,000) in 4 °C overnight, washed, and then incubated overnight at 4 °C with anti-rat HRP (Cell Signaling Technology, Cat#7077, 1:1,000). Detection used TSA.

For imaging, WISH samples were mounted in 80% glycerol supplemented with 10 mM Tris and 1 mM EDTA, pH = 7.5. Bright-field and WISH images were captured using a Canon EOS M50 mounted on a Zeiss Stemi 508 microscope. Samples for fluorescence imaging were mounted in scale solution (30% glycerol, 0.1% Triton X-100, 2 mg/mL sodium ascorbate, 4 M urea in PBS) and imaged on a Zeiss LSM 800 confocal microscope using 20× water immersion objective (N.A. = 1.0, working distance = 1.8 mm). All fluorescence images shown are maximum intensity projections and are representative of images taken in each condition. Cell counts (e.g., TUNEL^+^ or *ppib/c*^+^ cells) were quantified using the Fiji multi-point tool to mark cells through confocal z-stacks.

#### RNAi

dsRNA was prepared by *in vitro* transcription as described previously^53^. For RNAi through feeding, dsRNA was mixed with liver paste at a concentration of ∼150 ng/μL and fed to planarians every 4 d for 8 feedings for activin and activin receptor knockdowns, 3 feedings for *p38-1*, and 6 feedings for *fer3l-1, nkx2l, and ets-1*. For *act-2* RNAi in sexual planarians, animals were fed every 4 days for a total of 6 feedings. In all experiments, dsRNA matching ccdB and camR-containing insert of pJC53.2 was used as negative control. All primer sequences used in cloning for RNAi experiments are listed in **Table S2**.

#### Bulk RNAseq and data analysis

Total RNA was extracted using the RNeasy Micro Kit (Qiagen) from single animals. Libraries were prepared using ∼100 ng of total RNA as input using Universal Plus mRNA-Seq kit (Nugen) and sequenced on an Illumina Nextseq. Libraries were mapped to the dd_Smed_v6 transcriptome^58^ using bowtie2^59^. Counts from the same isotigs were summed, and pairwise differential expression analysis was performed using DEseq2^60^. Normalized read counts from DESeq2 were scaled to generate z-scores for heatmaps.

#### Planarian cell dissociation

A group of 10-15 mid-sized (∼5-7 mm in length) asexual planarians were finely minced with a razor blade, suspended in CMF (Ca/Mg-Free media: 480 mg/L NaH_2_PO_4_, 960 mg/L NaCl, 1.44 g/L KCl, 960 mg/L NaHCO_3_, 3.57 g/L HEPES, 0.24 g/L D-glucose, 1 g/L BSA, pH 7.4 in MilliQ H_2_O), and rocked for 15 min with gentle pipetting for 10 times every 3 min until visibly homogenized. Cells were centrifuged at 400 × g for 5 min and resuspended in 1 mL of fresh CMF, and serially filtered through 100, 70, 40 and 30-μm mesh strainers. After another spin, cells were resuspended in CMF supplemented with 1% BSA.

#### Cell lysis analysis ex vivo

For lysis assay, 50,000 cells were incubated overnight in 100 μL CMF supplemented with 1.5% FBS with control protein (h-CD8A) or ACT-2. Lysis was measured using Sony SH800 flow cytometry by the percentage of PI^+^ cells. We observed that ACT-2 concentrations required to induce cell lysis were higher than typical physiological levels. This is likely due to the recombinant protein’s lower biological activity compared to endogenous proteins, as well as potential underestimates of *in vivo* concentrations, as local protein concentrations can be significantly higher than tissue-wide averages.

To assess genotype specific responses, asexual and sexual cells were labeled with CFSE (Biolegend, Cat# 423801) or Celltrace Far Red (Invitrogen, Cat# 34564), respectively, and mixed at a 1:1 ratio with a total of 200,000 cells in 100 μL of CMF supplemented with 1.5% FBS.

For *E. coli* phagocytosis and dextran pinocytosis experiments, 50,000 total planarian cells were incubated overnight with 0.1 mg/mL pHrodo Green *E. coli* BioParticles (ThermoFisher Scientific, Cat# P35366) or 0.5 mg/mL Alexa Fluor 488-conjugated dextran (Invitrogen, Cat# D34682) in 100 μL CMF supplemented with 1.5% FBS and varying concentrations of ACT-2. Flow cytometry was used to quantify phagocytosis and pinocytosis by gating for live cells with high fluorescence intensity indicating the uptake of *E. coli* or dextran.

#### FACS

To isolate the ruptoblast population, dissociated cells were stained with Calcein Ultrablue AM (5 μM, AAT Bioquest, Cat# 21908) in CMF for 15 min at RT, washed, filtered, and incubated with PI (2 μg/mL) immediately before sorting on a Sony SH800 with a 130 μm chip. Cells were first gated for size using forward and back scattering, then based on Calcein Ultrablue and forward scattering (**Figure 3C and S5**). A total of ∼450,000 cells were sorted into CMF supplemented with 1% BSA, and 1.5% FBS for single-cell RNAseq experiment, ∼150,000 cells for time-lapse imaging, and ∼300,000 cells each population for Western blots.

#### Time-lapse imaging

Glass-bottom plates (Cellvis, Cat# P96-1-N) were coated with concanavalin A (50 μg/mL, Millipore, Cat# 11028-71-0) for 30 min. Approximately 50,000 cells were plated per well for 10 min before adding 20 μL of ACT-2 (10 μg/mL). Imaging began immediately on a Zeiss epifluorescence microscope using a EC Plan-Neofluar 10× objective (N.A = 0.3, working distance = 5.2 mm) or a LD Plan-Neofluar 63x objective (N.A = 0.75, working distance = 1.7 mm) at 2-12 frame/min. To label key cellular components, cells were incubated with fluorescent dyes including SiR-actin (1μM, Spirochrome, Cat# CY-SC001), SiR-DNA (1μM, Spirochrome, Cat# CY-SC007) and CellMask (1:1,000, Invitrogen, Cat#C10045). For imaging ruptoblast-mediated killing, P5 cells were labelled with CellTrace Violet (1:1,000, Invitrogen, Cat# C34571) and mixed 1:4 with unlabeled cells depleted of ruptoblasts (P1, P3, P4, P6). To quantify the timing and spatial distribution, images were binarized using OpenCV, and for each of the killed cell, the frame at which it first became PI^+^ was recorded along with its distance from the centroid of the adjacent ruptoblast.

For bacterial killing experiments, 96-well plates were coated with Poly-L-lysine (Cultrex, Cat#3438-100-01) for 30 min. 3 μL of *E. coli* (MG1655 attB::pproC-msfGFP, OD ∼1.2)^61^ were diluted in 60 µL of CMF and plated for 30 min. About 50,000 P5 cells in 100 μL of CMF with PI (2 μg/mL) were added and co-incubated 20 min prior to introducing 20 μL of ACT-2 (10 μg/mL) and proceeding to imaging. To quantify GFP^+^ and PI^+^ bacterial fractions, images were binarized using OpenCV and small debris were excluded by applying a minimum area threshold of >10 pixels.

To visualize calcium dynamics during ruptosis, cells were incubated with Fluo-4 AM (1:1000, Invitrogen, Cat# F14217). For ruptosis perturbation experiments, 25,000 P5 cells were incubated under the following conditions prior to activation with ACT-2 (10 μg/mL): 250 mM sucrose (400 mOsm/L) in CMF or 125 mM sucrose (250 mOsm/L) in CMF for 45 min; 10 μM blebbistatin (STEMCELL Technologies, Cat# 72402) or 10 μM latrunculin A (Invitrogen, Cat# L12370) for 1 hr; 10 μM jasplakinolide (Invitrogen, Cat# J7473) for 45 min; and 20 μM BAPTA-AM (Invitrogen, Cat# B1205) for 1 hr.

To quantify granule release during ruptosis, time-lapse images were binarized using OpenCV, and the number of granules was manually counted using Fiji’s multipoint tool. Counts were averaged over five consecutive frames corresponding to the period of maximum granule dispersal. To measure changes in cell area following perturbation, images were first processed in ImageJ with a convolution filter (e.g., Gaussian blur) to enhance contrast and suppress background noise. The resulting images were then binarized, and cell areas were measured and tracked over time.

#### Single-cell RNAseq and data analysis

Following FACS, 150,000 or 300,000 P5 cells were incubated for 5 min in CMF supplemented with 1% BSA containing either control protein (h-CD8A, 10 μg/mL) or ACT-2 (10 μg/mL), respectively. Cells were then centrifuged at 400 × g, resuspended in fresh CMF with the same protein for another 5 min, centrifuged again, and finally resuspended in CMF at ∼1,000 cell/μL. Libraries were prepared using a 10x Genomics Chromium Controller with Chromium single cell v3.1 library/Gel Bead Kit. Amplified cDNA libraries were quantified with a bioanalyzer and sequenced on an Illumina Novaseq S4 platform, generating in mean coverage of ∼25,000 read pairs per cell.

UMI-tools^62^ and cutadapt^63^ were used for barcode tagging and adapter trimming, respectively. Reads were aligned to dd_Smed_v6 transcriptome using bowtie2 with “–sensitive” parameters. Ambient RNA contamination was removed using SoupX^64^ with default parameters. After removing low quality cells with fewer than 200 genes detected, we captured 1,500 genes and 4,000 UMI on average for the h-CD8A-treated control sample and 1,300 genes and 3,200 UMI for the ACT-2-treated sample. Counts were normalized such that each cell’s total matched the median library size, and log_2_-transformed after adding a pseudo count of 1. Harmony^65^ was used to integrate control and ACT-2 samples and SAM^66^ was used for manifold reduction with default parameters, except for weight_mode = rms. Clusters were annotated based on known marker genes^23^. Depletion factor was calculated as the ratio of relative abundance of each cluster in ACT-2 vs. control samples.

To identify ruptoblasts in the full planarian cell atlas and examine the specificity of their marker genes, we reanalyzed the published single-cell data^23^ using the same pipeline, filtering out cells with <500 genes.

#### Pseudomonas infection and commensal bacterial load quantification

*Pseudomonas* isolated from the planarian^25^ was inoculated in LB at 30 °C for 24 hr. OD600 was measured and converted to CFU/mL. Bacterial pellets were washed and resuspended in IO salts. Asexual animals (∼3-4 mm in length) were washed several times before transferring to petri dishes containing bacteria. This process was repeated every 3 days.

Planarians (∼3-4 mm in length) were homogenized with 6 animals in 100 μL of milli-Q using a pestle, and the entire 100 μL was plated on LB agar without antibiotics. Plates were incubated in the dark at RT for 24 hr. A water-only control verified no contamination.

#### Mammalian cell killing

HEK293 cells were cultured in a controlled humidified incubator at 37 °C and 5% CO_2_ in DMEM (Gibco, Cat# 10569069) supplemented with GlutaMAX, sodium pyruvate, 10% FBS (Omega Scientific, Cat# 20014T), and 1% Penicillin-Streptomycin-Glutamine (Gibco, 10378016) and seeded into black, glass-bottom 96-well plates (Cellvis, Cat# P96-1-N) at a density of 20,000 cells per well, 24 hr prior to imaging. RAW264.7 cells were maintained in high-glucose DMEM (Gibco, Cat# 11965092) supplemented with 10% FBS and 1% Penicillin-Streptomycin-Glutamine, and seeded at 30,000 cells per well 16 hr prior to imaging. Following removal of the culture medium, ∼25,000 P5 cells suspended in PBS were added to each well and allowed to settle for 5 min. 20 μL of ACT-2 (10 μg/mL) was then added and imaging was initiated immediately.

To measure caspase-1 activation in RAW cells, ∼25,000 P5 cells resuspended in 100 μL PBS were mixed with ACT-2 protein (10 μg/mL) and FAM-YVAD-FMK caspase-1 inhibitor reagent (1:30 dilution of 30X FLICA stock in PBS; Pyroptosis/Caspase-1 Assay Kit, ImmunoChemistry Technologies, Cat# 9145). The mixture was added to 96-well plate containing pre-seeded 30,000 RAW cells per well and incubated for 5 min at RT, followed by 1 hr incubation at 37 °C in a humidified incubator with 5% CO₂. After incubation, cells were washed three times with the provided wash buffer, stained with propidium iodide (PI, 2 μg/mL) for 10 min, and immediately imaged.

To access the cytotoxic potency of the supernatant, 50,000 P5 cells resuspended in 50 μL PBS were incubated with ACT-2 (10 μg/mL) for 5 min, followed by centrifugation at 500 × g for 5 min. The resulting supernatant (∼ 48 μL) was collected and added to pre-seeded HEK293 cells, and imaging was initiated immediately. For mechanical lysis of ruptoblasts, 100,000 P5 cells resuspended in 100 μL PBS were homogenized using a handheld pestle homogenizer for ∼ 5 min, followed by centrifugation at 500 ×g for 5 min. The resulting supernatant (∼50 uL) was then collected and applied to HEK293 cells.

For DNase treatment, the supernatant was incubated with DNase I (100 U/mL, Promega, Cat# M6101) at 37 °C for 5 min prior to adding to the cells. To determine the molecular size range of the active cytotoxic agents, 100,000 P5 cells resuspended in 100 μL PBS were incubated with ACT-2 (10 μg/mL), followed by centrifugation at 500 × g for 5 min. The collected supernatant was passed through 30 kDa (Amicon, Cat# UFC503008) or 100 kDa (Amicon, Cat# UFC510008) molecular weight cutoff (MWCO) centrifugal filters by spinning at 12,000 × g for 5 min and 8,000 × g for 5 min, respectively. The resulting filtrates were then applied to HEK293 cells.

## Supplemental Tables

**Table S1.**
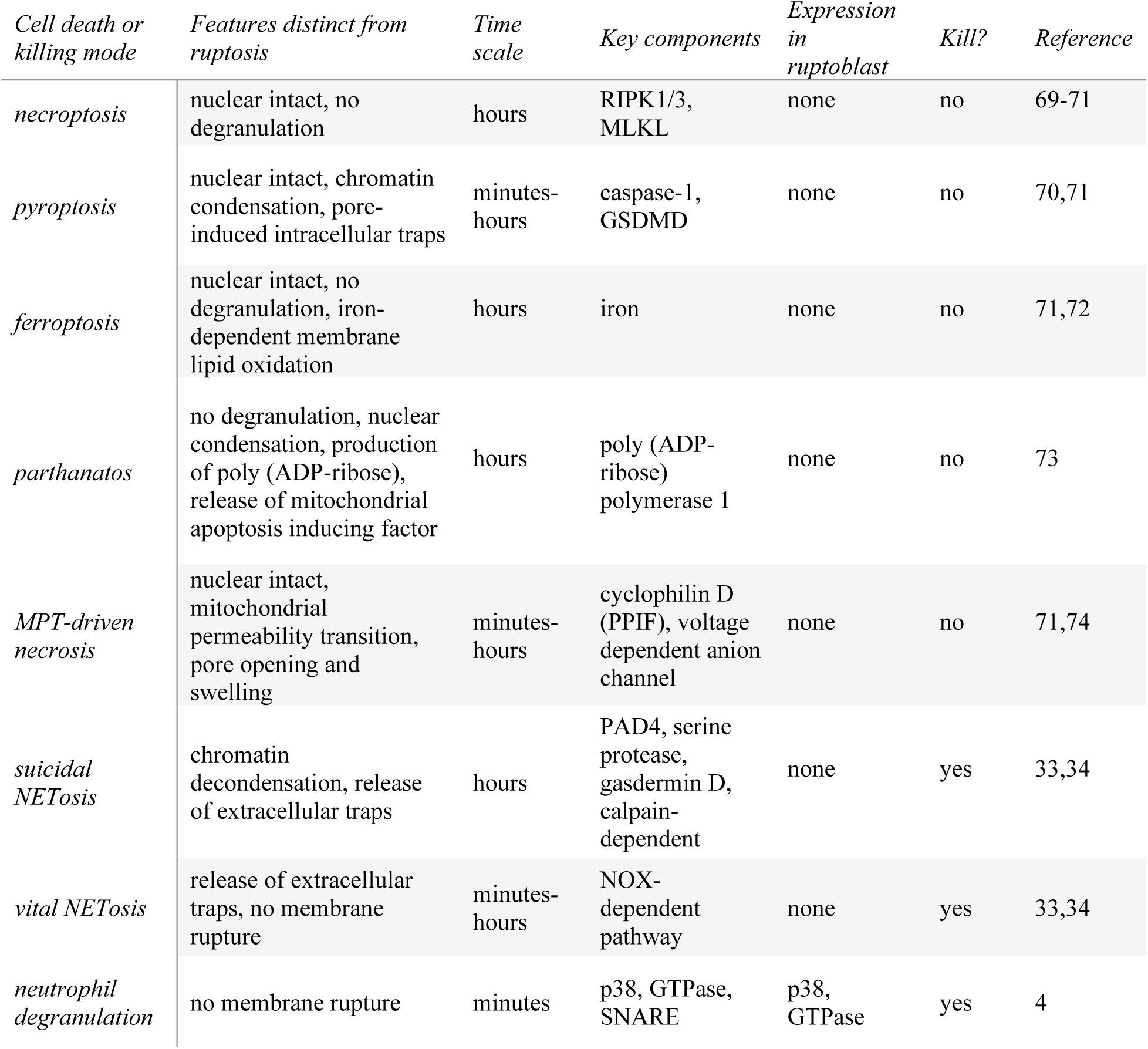
Summary of previously characterized cell death processes, outlining their key cellular events, timescales and molecular regulators that are different from ruptosis.

**Table S2.** List of gene IDs associated with planarian genes used throughout the study along with primer sequences used in cloning for in-situ hybridization and RNAi experiments.

## Supplemental Videos

**Video S1.** Brightfield videos of total planarian cells incubated with 10 μg/mL of control proteins (h-CD8A or h-ACTB) or ACT-2 protein, along with videos of sorted subpopulations (P1, P3–P6) incubated with 10 μg/mL ACT-2 proteins. Only P5 responds to ACT-2. As a comparison, P5 incubated with 10 μg/mL control (h-CD8A) protein is shown to demonstrate the absence of lysis. Proteins are added at 0 s. Time stamp: minute:second.

**Video S2.** Brightfield and fluorescence videos of ruptoblasts undergoing ruptosis upon activation with 10 μg/mL of ACT-2 along with videos of ruptoblasts pretreated with 250 mM sucrose for 45 min before adding ACT-2. Ruptoblasts are labelled with CellMask (yellow) for plasma membrane, SiR-DNA (blue) for nucleus, and SiR-actin (red) for actin cytoskeleton. ACT-2 is added at 0 s. Time stamp: minute:second.

**Video S3.** Time-lapse video of ruptoblasts labeled with CellTrace Violet (blue) killing nearby unlabeled planarian cells, as indicated by PI uptake (red). ACT-2 is added at 90 s. Time stamp: minute:second.

**Video S4.** Time-lapse video showing ruptoblast killing nearby *E. coli* upon activation by 10 μg/mL ACT-2, as indicated by the loss of cytoplasmic GFP (green) followed by PI uptake (red). As a control, P4 cells incubated with *E. coli* along with 10 μg/mL ACT-2 is shown afterwards, demonstrating the absence of bacterial lysis. ACT-2 is added at 0 s. Time stamp: minute:second.

**Video S5.** Time-lapse videos showing ruptoblasts killing HEK293 and RAW264.7 cells upon activation by 10 μg/mL ACT-2, as indicated by PI uptake (red), followed by videos showing HEK293 cells incubated with supernatant collected from ruptoblasts activated with ACT-2 or supernatant obtained by mechanical lysis of ruptoblasts without ACT-2 activation. ACT-2 is added at 0 s. Time stamp: minute:second.

**Video S6.** Time-lapse videos of calcium dynamics during ruptosis followed by videos of ruptoblasts undergoing ruptosis under different perturbation conditions. Ruptoblasts were either pretreated with 10 μM blebbistatin or 10 μM Lat A for 1 hr, or 10 μM JAS for 45 min, or 20 μM BAPTA-AM for 1 hr, before activation by 10 μg/mL ACT-2. Cells were labeled with CellMask (yellow), SiR-DNA (blue), and SiR-actin. ACT-2 is added at 0 s. Time stamp: minute:second.

## Notes

### Competing Interest Statement

The authors have declared no competing interest.

### Summary of Updates

We have added two new result section in which we show cross-species cytotoxicity of ruptoblasts as well as cell biology experiments in which we show the explosive nature of ruptoblast is regulated by calcium signaling and cytoskeleton dynamics.

